# JNK integrates immune and stress signals to balance apoptosis and proliferation in airway progenitors

**DOI:** 10.1101/2025.10.30.685523

**Authors:** Leizhi Shi, Xiao Niu, Kunyu Zhang, Karina Stein, Holger Heine, Jörg U. Hammel, Iris Bruchhaus, Susanne Krauss-Etschmann, Judith Bossen, Thomas Roeder

## Abstract

Chronic inflammation disrupts epithelial regeneration, yet how immune signaling reprograms progenitor fate remains unclear. Using the *Drosophila* airway as a model of epithelial remodeling, we identify the c-Jun N-terminal kinase (JNK) pathway as a central integrator of immune and stress cues that balances apoptosis and compensatory proliferation in airway progenitors. Persistent activation of the innate immune IMD pathway induces simultaneous cell death and proliferation through a non-canonical route that bypasses NF-κB/Relish and instead engages JNK. Downstream, distinct transcriptional modules orchestrate these divergent fates: Foxo and AP-1 drive apoptosis, whereas the ETS factor Ets21C mediates proliferation. Genetic inhibition of JNK or its effectors restores progenitor homeostasis, while constitutive activation recapitulates inflammation-induced tissue remodeling. These responses are cell-autonomous, revealing that airway progenitors actively interpret immune and stress signals to determine their fate. Collectively, our findings uncover a modular signaling architecture that links inflammation to regeneration and highlight conserved JNK-dependent transcriptional programs as potential therapeutic targets to prevent progenitor exhaustion in chronic airway disease.

## Introduction

The lungs are continually exposed to a diverse array of environmental insults, including airborne pathogens, pollutants, allergens, and toxic particulates. To counter these challenges, a robust and localized immune defense is essential. The airway epithelium not only forms a physical barrier but also plays a central role in immune surveillance, coordinating the recruitment of immune cells and initiating inflammatory responses [1, 2]. While acute inflammation is vital for pathogen clearance, chronic inflammation drives lung pathology and underlies diseases such as asthma and chronic obstructive pulmonary disease (COPD) [3–8].

The airway epithelial barrier is intrinsically dynamic [8, 9] and comprises a mosaic of functionally specialized cell types [10]. Differentiated epithelial cells actively sense environmental cues and secrete antimicrobial peptides, chemokines, and cytokines to modulate immune responses [10]. In contrast, progenitor cells—such as basal cells in the proximal airway and alveolar type 2 (AT2) cells in the distal lung—are primarily responsible for maintaining epithelial integrity and orchestrating repair following injury [11].

Although traditionally viewed as quiescent and uninvolved during inflammation, progenitor cells are increasingly recognized as active participants in the immune milieu. Emerging evidence suggests that these cells can sense inflammatory cues and adjust their behavior accordingly [12]. Chronic inflammatory stress can impair progenitor cell function, reducing their regenerative capacity—a phenomenon linked to both lung aging and the concept of “inflammaging” [13, 14]. Interestingly, the effects of inflammation on progenitor activity are context-dependent: immune signals can either enhance or suppress progenitor proliferation and differentiation, depending on the timing, duration, and nature of the insult [15, 16]. This duality is particularly evident in the lung, where basal and AT2 cells exhibit highly plastic, stimulus-specific responses to immune activation [17–19].

Despite their essential roles in maintaining tissue homeostasis, the molecular mechanisms by which inflammation regulates the fate of progenitor cells remain poorly defined. Beyond their regenerative functions, progenitor cells may also serve as long-lived cellular recorders of prior immune events [20]. Dissecting how inflammatory signals shape their developmental trajectories is thus critical to understanding both tissue regeneration and the progression of chronic lung disease [21, 22].

To address this knowledge gap, we employed a genetically tractable model that recapitulates key features of mammalian respiratory progenitor biology: the tracheoblasts of the spiracular branch (SB) in *Drosophila melanogaster* [23–26]. These cells share functional parallels with vertebrate airway progenitors, providing a robust system for *in vivo* manipulation and lineage tracing. Our investigation focused on the tracheoblasts of metameres 4 and 5, which contribute to the development of the adult tracheal system. We demonstrate that these progenitor cells mount a robust and context-specific response to immune stimulation, actively reprogramming their developmental fate in response to inflammatory cues. Our findings reveal that airway progenitor cells function not merely as repair agents but as direct interpreters of immune and stress signals. These results uncover conserved principles of epithelial plasticity that govern regeneration during immune challenge.

## Results

The respiratory organ of *Drosophila*, the trachea, is organized as a complex, blind-ending network of tubes (Fig. 1A), where also progenitor cells that make the adult tracheal system are part of (Fig. 1B). Bacterial infection of this *Drosophila* respiratory system induces a strong epithelial antimicrobial response, marked by the upregulation of multiple antimicrobial peptide (AMP) genes□[27]. To visualize this response, we infected the airways of a reporter strain in which GFP expression is driven by the native *drosomycin* promoter, using the insect pathogen *Pectobacterium carotovorum* [28, 29] (Fig. 1C-E).

**Figure 1.**
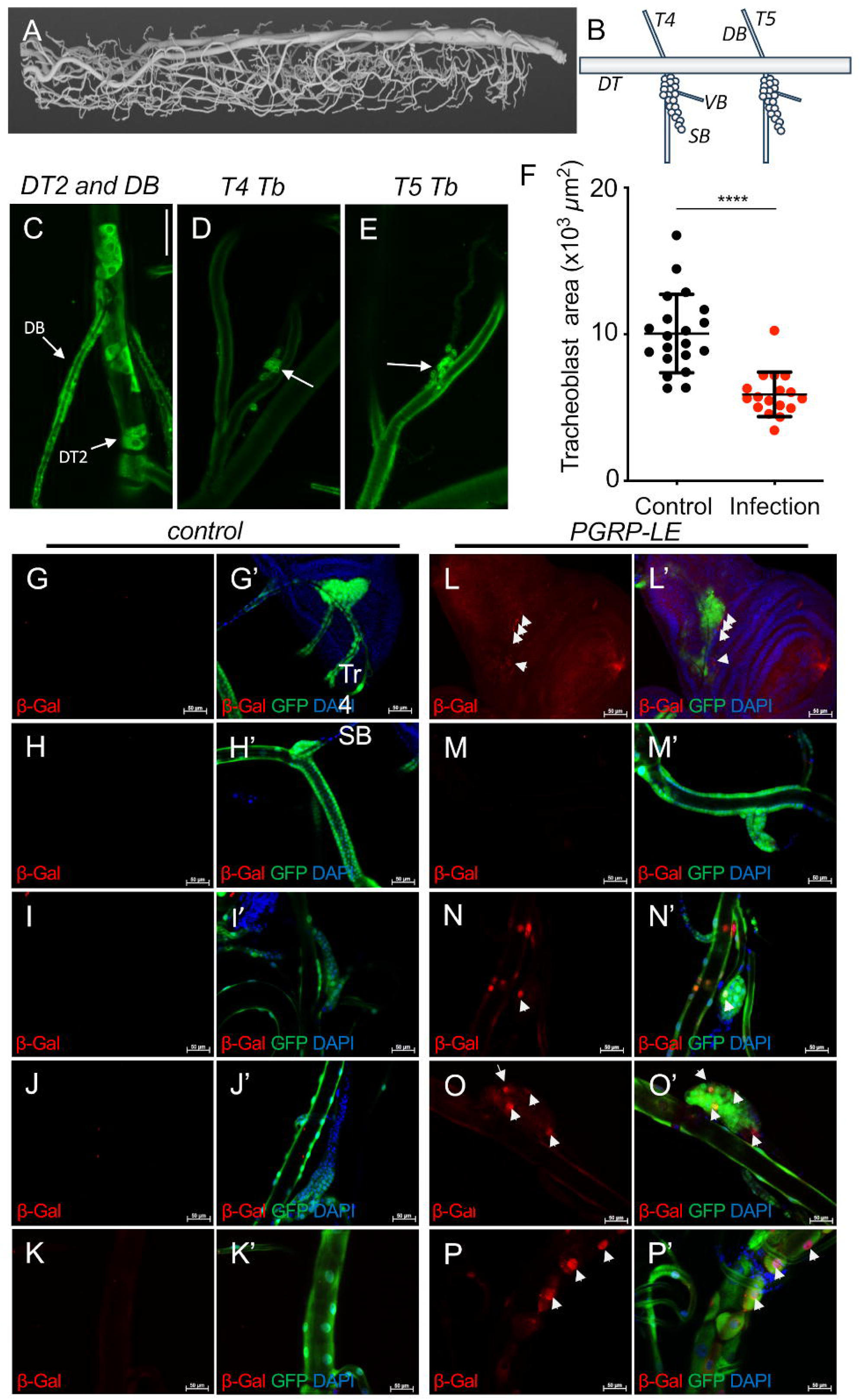
Infection induces expression of AMP genes in progenitor cells of the larval trachea. A reconstruction of the tracheal system of L3-stage *Drosophila* larvae using µCT analysis (A). Schematic description of the metameres T4 and T5 major parts of the tracheal system, highlighting the SB tracheoblasts (B). *Drosophila* (*Drs-GFP; Dipt.-lacZ*) L3 larva subjected to an infection protocol targeting the trachea (C-F). Dissected trachea showing GFP-positive cells in the Dorsal Branch (DB) and the DT2 area (C), the T4 Tracheoblast (D), and the T5 tracheoblast (E). Effect of an airway infection on the size of the tracheoblasts (F). Dissected trachea from control larvae (E-I, E’-I’: *btl-Gal4, UAS-GFP; tub-Gal80 [ts] > w^1118^*), PGRP-LE overexpressing larvae (J-N, J’-N’: *btl-Gal4, UAS-GFP; tub-Gal80 [ts] > PGRP-LE (DDI (Drs-GFP; Dipt.-lacZ): UAS-Flag-PGRP-LE)*. Isolated tracheae were stained with β-galactosidase antibody (red). Tracheal tissue was simultaneously stained with anti-GFP (green) and DAPI (blue (G’-P’). β-galactosidase-positive cells are highlighted by white arrows. (G, G’, L, L’) ASP cells. (H, H’, J, J’) DT2 cells. (I, I’, N, N’) T4 tracheoblasts. (J, J’, O, O’) T5 tracheoblasts. (K, K’, P, P’) Dorsal trunk. Scale = 50 µm. **** p < 0.001.

**Figure 2.**
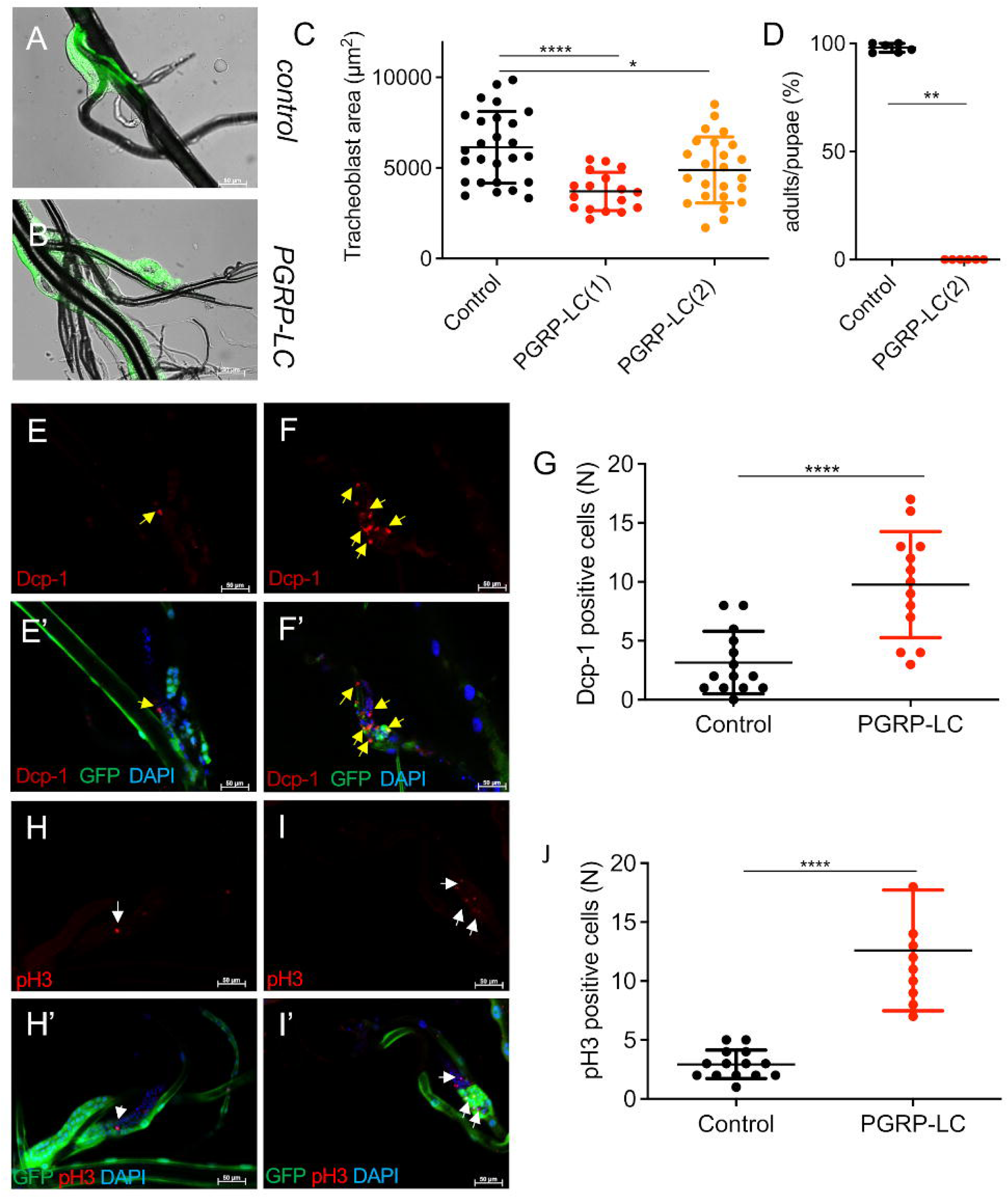
Constitutive activation of the IMD pathway in the airways induces animal death and shrinkage of the tracheoblast area. (A) Dissected tracheae from *btl-Gal4, UAS-GFP; tub-Gal80 [ts] > w^1118^* control larvae. *Btl-Gal4, UAS-GFP; tub-Gal80 [ts] > UAS-PGRP-LC(1)* larvae with ectopic expression of *PGRP-LC* in the trachea (B). (C) Quantitative analysis of the area of tracheoblasts. Eclosion probability of animals subjected to PGRP-LC overexpression in the trachea. The proportion of hatched adults per pupae is shown (D). Dissected tracheae from control larvae (E, E’, H, H’; *btl-Gal4, UAS-GFP; tub-Gal80 [ts] > w^1118^*), PGRP-LC larvae (F, F’, I, I’; *btl-Gal4, UAS-GFP; tub-Gal80 [ts] > UAS-PGRP-LC*). (E, E’, F, F’) were stained with the Dcp-1 antibody (red). (H, H’, I, I’) were stained with the anti-pH3 antibody (red). (E’, F’, H’. I’) Tissue of trachea stained GFP in green, nuclear DAPI in blue. Yellow arrows (pH3 positive cells, white arrows) highlight Dcp-1 positive cells. (G) Statistical analysis of the dcp-1 positive cells of T5 tracheoblasts. (J) Statistical analysis of pH3-positive cells of T5 tracheoblasts. Scale bar = 50 µm. Statistical significance was evaluated by Mann–Whitney test, ***p < 0.001, ****p < 0.0001. n≥10. Scale bar = 50 µm. Statistical significance was evaluated by Mann–Whitney test, *p < 0.05, **p < 0.01, ****p < 0.0001. n≥10.

In uninfected controls, no GFP signal was detected in the tracheal system. In contrast, infected flies showed robust *drosomycin* activation in the airway epithelium as well as in tracheal progenitor cells, including DT2 cells in the dorsal trunk of the second metamere and dorsal branch (DB) cells (Fig. 1C), T4 and T5 spiracular branch (SB) tracheoblasts (Fig. 1D, E). Notably, in SB tracheoblasts, the tracheal area covered was significantly reduced following infection (Fig. 1F).

To investigate the mechanisms underlying these morphological changes, we employed a genetically controlled activation system. Specifically, we stimulated the IMD immune pathway in airway tissues by targeted overexpression of the pattern-recognition receptors *PGRP-LE* or *PGRP-LC*. To ensure activation in all immunocompetent tracheal cells, including SB tracheoblasts, *PGRP-LE* was expressed under the *btl*-Gal4 driver (31, 32). Secreted PGRP-LE engages PGRP-LC–expressing cells connected to the tracheal lumen.

To monitor IMD pathway activation, we co-expressed two reporters—*Diptericin-lacZ* (β-Gal, red) and *Drosomycin*-GFP (green)—with *PGRP-LE* (Fig. 1G–P’). In the absence of ectopic *PGRP-LE*, *Diptericin* expression was undetectable in any tracheal compartments, including the air sac primordia (ASP; Fig. 1G, G’), DT2 tracheoblasts (Fig. 1H, H’), T4 (Fig. 1I, I’), and T5 SB tracheoblasts (Fig. 1J, J’), and the dorsal trunk (Fig. 1K, K’).

By contrast, *PGRP-LE* overexpression strongly induced *Diptericin*-positive cells in multiple regions, including the ASP (Fig. 1L, L’), T4 and T5 SB tracheoblasts (Fig. 1N, N’; O, O’), and dorsal trunk (Fig. 1P, P’). Interestingly, activation was not observed in DT2 tracheoblasts (Fig. 1M, M’). Although SB tracheoblasts are present in all tracheal segments, we focused subsequent experiments on the T4 and T5 SB tracheoblasts due to their central role in adult tracheal system formation.

### Constitutive activation of IMD signaling in the Drosophila airway epithelium leads to marked epithelial thickening and a significant reduction in SB tracheoblast size

Our experimental design ensured cell-autonomous IMD pathway activation, a crucial element for mechanistic analysis. We achieved this by using membrane-bound PGRP-LC to restrict pathway induction specifically to PGRP-LC-expressing cells. Temporal control was achieved via a Gal80^ts^ system, which represses at 19 °C and is induced upon a temperature shift [30].

Consistent with previous findings [31], PGRP-LC overexpression in the larval tracheal system triggered pronounced epithelial thickening. Upon inducing PGRP-LC expression in early third instar (L3) larvae for 2 days (shift from 18 °C to 29 °C), we observed a significant increase in tracheal epithelial thickness relative to controls (quantified in the T9 segment; see Supplementary Figure 1A–D’).

Focusing on SB tracheoblasts, we report here the results for T5 SB tracheoblasts. To dissect downstream pathway elements, two independent PGRP-LC alleles on separate chromosomes were employed: PGRP-LC(1) on chromosome II and PGRP-LC(2) on chromosome III (Fig. 2C). This strategy enabled simultaneous genetic manipulation of candidate downstream genes.

Activation of the IMD pathway via PGRP-LC overexpression resulted in a significant reduction in SB tracheoblast area in the T5 segment (control: 6,139 ± 387.9 µm²; PGRP-LC(1): 3,709 ± 255.6 µm², p < 0.0001; PGRP-LC(2): 4,892 ± 371.4 µm², p = 0.0343; Fig. 2C), and the SB tracheoblasts repositioned from the dorsal trunk to the secondary branches following induction (Fig. 2A,B).

To test effects on airway development, we induced PGRP-LC expression for 6 days at the repressive temperature (19 °C), followed by transfer to the permissive condition (29 °C) using Gal80^ts. Developmental outcomes were assessed by calculating the adult-to-pupa ratio. While most animals reached the pupal stage, none expressing ectopic PGRP-LC survived to adulthood (Fig. 2D).

### Constitutive IMD activation induces apoptosis and compensatory proliferation in SB tracheoblasts

To determine whether the reduction in SB tracheoblast area following tracheal PGRP-LC overexpression results from increased apoptosis, we stained for the apoptotic marker Dcp-1. Tracheae with PGRP-LC overexpression exhibited a significant increase in Dcp-1–positive cells within the SB branches (Fig. 2E, F; yellow arrows), with nearly a threefold elevation in T5 SB tracheoblasts (9.77 ± 1.25) compared to controls (3.14 ± 0.71; p < 0.0001; Fig. 2G).

In controls, no Dcp-1 staining was detected in dorsal trunks (Supplementary Fig. 2), while only sporadic Dcp-1–positive cells appeared in dorsal trunks upon PGRP-LC overexpression. Apoptotic vesicles were readily observed in affected cells (Supplementary Fig. 2B, B’, white arrows). Extended PGRP-LC induction for 2 days led to an even greater number of Dcp-1–positive cells (Supplementary Fig. 2C, C’, yellow arrows).

To assess whether compensatory cell proliferation occurred alongside increased cell death, we used anti-phospho-histone H3 (pH3) antibody staining to detect mitotically active SB tracheoblasts (Fig. 2H–I’; white arrows). The number of pH3-positive cells in PGRP-LC–activated T5 SB tracheoblasts was significantly higher than in controls (Fig. 2J).

### Downstream components that mediate the effects of IMD pathway activation on the SB tracheoblasts

TAK1, an integral part of the IMD signaling pathway, interacts with Tab2 to form a hub that connects the IMD and JNK signaling pathways [31, 32]. Therefore, we examined the effect of reducing TAK1 activity by co-expressing a dominant-negative TAK1 allele (dTAK1^DN^) concurrently with *PGRP-LC* overexpression (Figure 3A–D). We observed a substantial increase in SB tracheoblast sizes (exceeding even that of the wild type) in flies overexpressing *PGRP-LC* concurrently with dTAK1^DN^ in the tracheal system when compared with *PGRP-LC* overexpression only (Figure 3A-D, data for the T4 tracheoblasts are shown in Supplementary Figure 3).

To determine whether activating Relish, the canonical NF-κB protein of the IMD pathway, is sufficient to phenocopy the effects of *PGRP-LC*-induced IMD pathway activation, we used *Rel-V5* (Figure 3E), an inactive cleavage product of Relish, and the constitutively active allele *Rel-D* (Figure 3F) [33]. Overexpression of *Rel-D* did not phenocopy the effects of IMD pathway activation on tracheoblast size; instead, a slight increase in size was observed (Figure 3G), which shows that Relish activation is not sufficient to induce the small tracheoblast area phenotype (data for the T4 tracheoblasts are shown in Supplementary Figure 3).

**Figure 3.**
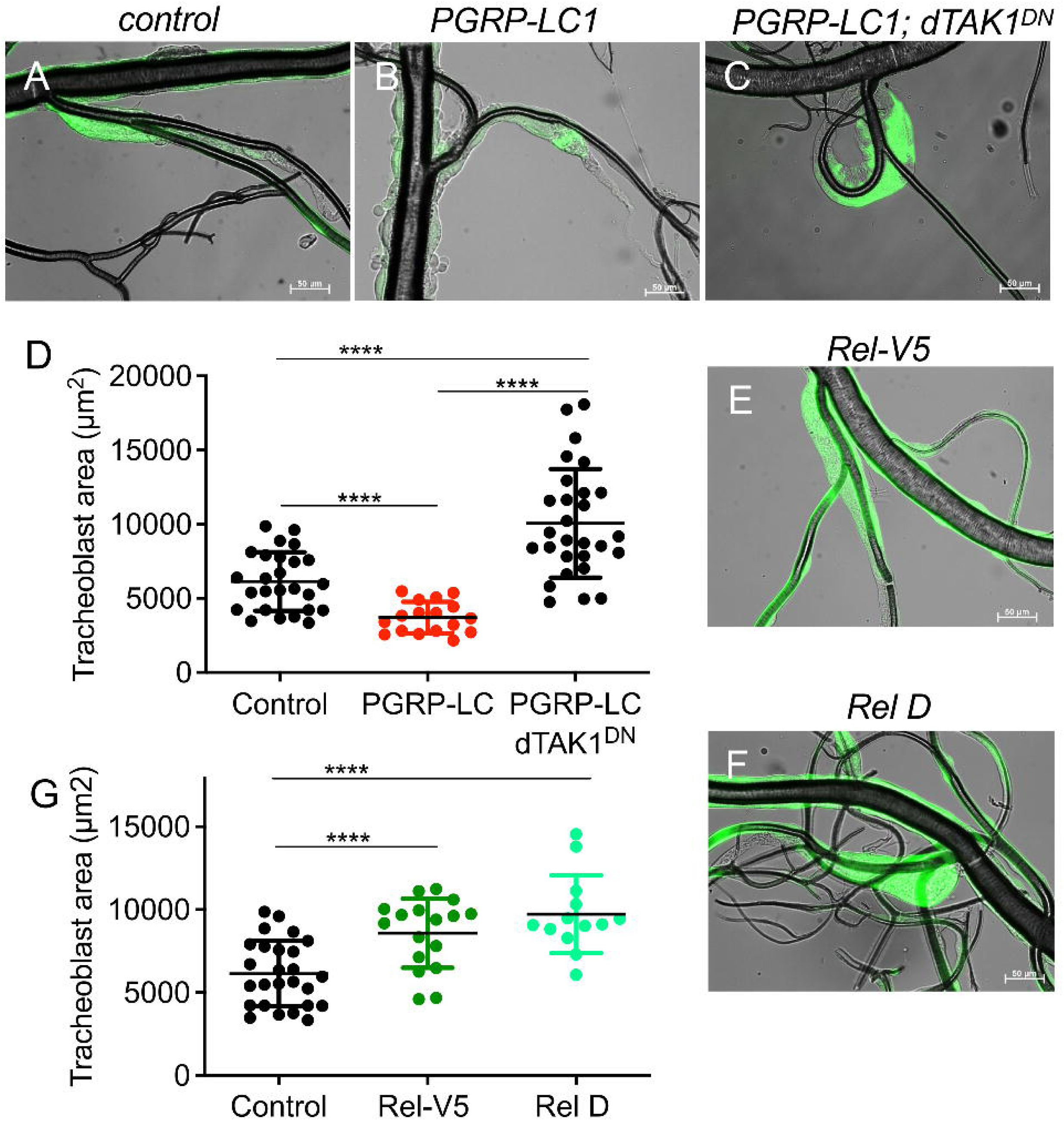
IMD-induced reduction in the tracheoblast area does not depend on the canonical Relish signaling pathway downstream of TAK1. Dissected tracheae from *btl-Gal4, UAS-GFP; tub-Gal80 [ts] > w^1118^* control larvae (A). Dissected tracheae from *Btl-Gal4, UAS-GFP; tub-Gal80 [ts] > UAS-PGRP-LC(1)* (B). Dissected tracheae from *Btl-Gal4, UAS-GFP; tub-Gal80 [ts] > UAS-PGRP-LC(1); UAS-dTAK1^DN^* (C). (D) Quantitative analysis of the area of T5 tracheoblasts. (E) Dissected tracheae from *btl-Gal4, UAS-GFP; tub-Gal80 [ts] > UAS-Rel-V5*. (F) Dissected tracheae from *btl-Gal4, UAS-GFP; tub-Gal80 [ts] > UAS-Rel-D. (G)* Quantitative analysis of the area of T5 tracheoblasts. Images of control w^1118^ for E and F are the same as A and are not shown again. Scale bar = 50 µm. Statistical significance was evaluated by Mann–Whitney test, ns, not significant, *p < 0.05, **p < 0.01, ***p < 0.001, ****p < 0.0001. n≥10.

### Reduced SB Tracheoblast Size Upon IMD Activation Requires JNK Signaling

Since Relish activation alone does not reproduce the IMD-induced reduction in tracheoblast size, we tested whether this phenotype is mediated through JNK signaling downstream of dTak1, in line with our previous finding that IMD activation at the dorsal branch level diverts into the JNK pathway to regulate epithelial thickness [34].

To inhibit JNK activity, we co-expressed a dominant-negative basket (bsk^DN^) allele— encoding *Drosophila* JNK—together with PGRP-LC in the tracheal system (Fig. 4). Blocking JNK signaling fully rescued the PGRP-LC–induced reduction in T4 and T5 SB tracheoblast size, restoring areas to control levels and even slightly exceeding them (Fig. 4A–D). This indicates that the size phenotype is entirely JNK-dependent. Because basket inhibition completely abolished the PGRP-LC-induced size reduction, we examined whether it also suppressed the two associated molecular phenotypes—apoptosis and proliferation. Indeed, blocking JNK restored Dcp-1– positive apoptotic cell counts in T4 and T5 SB tracheoblasts to near-control levels (Fig. 4E–H, Supplementary Fig. 4). Similarly, the PGRP-LC–induced increase in pH3-positive mitotic cells (Fig. 4J, J’, L) was fully rescued to baseline levels upon bsk^DN^ expression (Fig. 4K, K’, L).

Next, we tested whether direct activation of the canonical JNK pathway could phenocopy this effect (Fig. 4M). Overexpression of the constitutively active TNF receptor Grindelwald (Grnd^CA^), either in all tracheal cells (*btl*-Gal4) or specifically in tracheoblasts (*Ci*-Gal4), led to a highly significant reduction in tracheoblast area, indistinguishable from that caused by immune activation via PGRP-LC (Fig. 4N, O). In summary, activation of the JNK pathway downstream of dTak1 is both necessary and sufficient to mediate IMD-induced reductions in SB tracheoblast size as well as the concomitant increases in apoptosis and proliferation.

**Figure 4.**
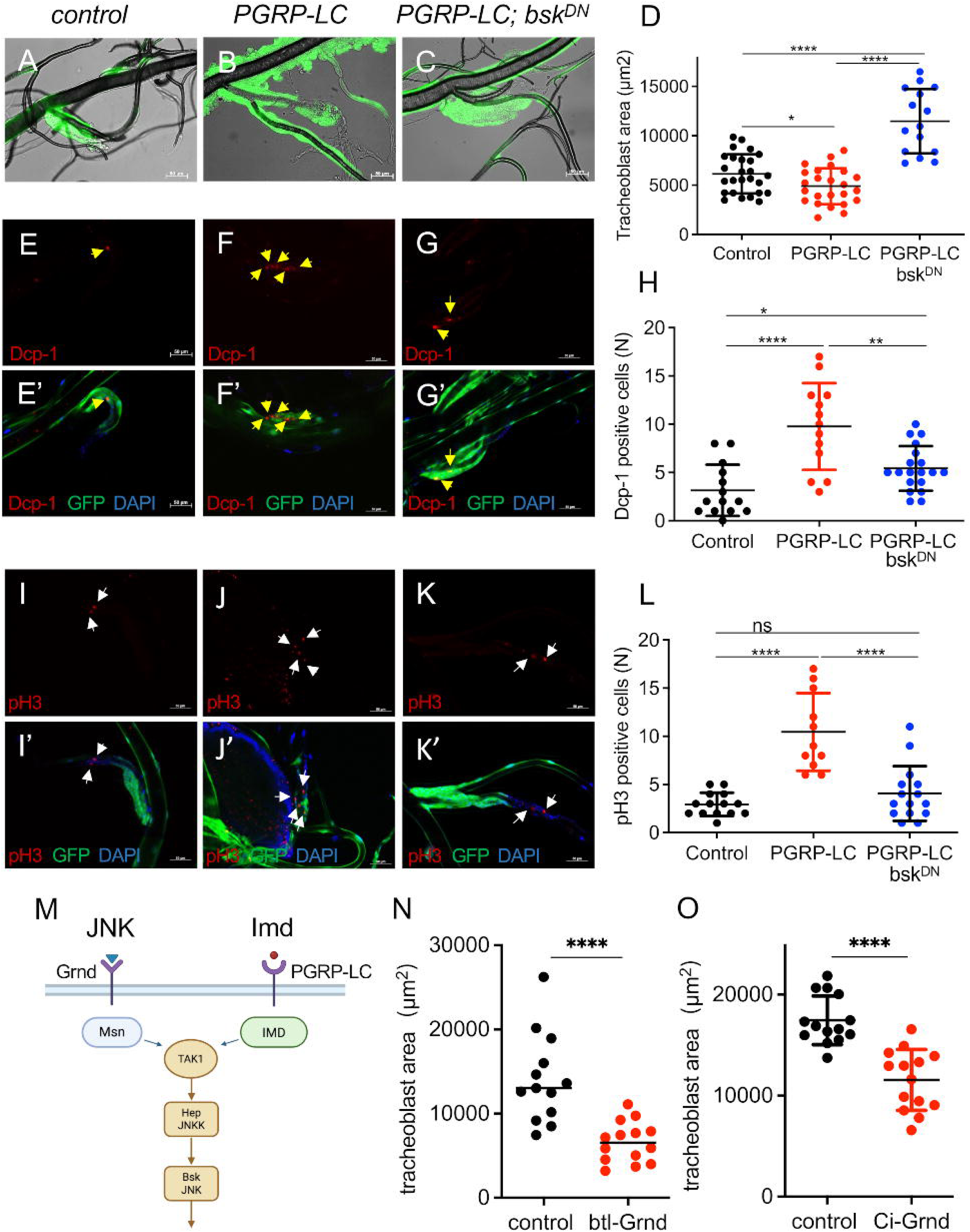
The effects on T5 tracheoblasts caused by activation of IMD signaling are rescued by JNK (bsk) silencing. (A, I, I’) Dissected tracheae from *btl-Gal4, UAS-GFP; tub-Gal80 [ts] > w^1118^*. (B, F, F’) Tracheae from *Btl-Gal4, UAS-GFP; tub-Gal80 [ts] > UAS-PGRP-LC(2)*. (C, G, G’) Dissected tracheae from *Btl-Gal4, UAS-GFP; tub-Gal80 [ts] > UAS-PGRP-LC(2); UAS-Bsk^DN^*. (D) Quantitative analysis of the area of T5 tracheoblasts. (E-H) Effects of PGRP-LC activation and of concurrent *bsk* silencing on pH3-positive cell numbers. (I-L) Effects of PGRP-LC activation and of concurrent *bsk* silencing on dcp-1-positive cell numbers. (H) Quantitative analysis of the dcp1-positive cells per tracheoblast. (L) Quantitative analysis of the pH3-positive cells per tracheoblast. Arrows mark positive cells. Scale bar = 50 µm. Statistical significance was evaluated by Mann–Whitney test, ns, not significant, *p < 0.05, **p < 0.01, ***p < 0.001, ****p < 0.0001, n≥10.

**Figure 5.**
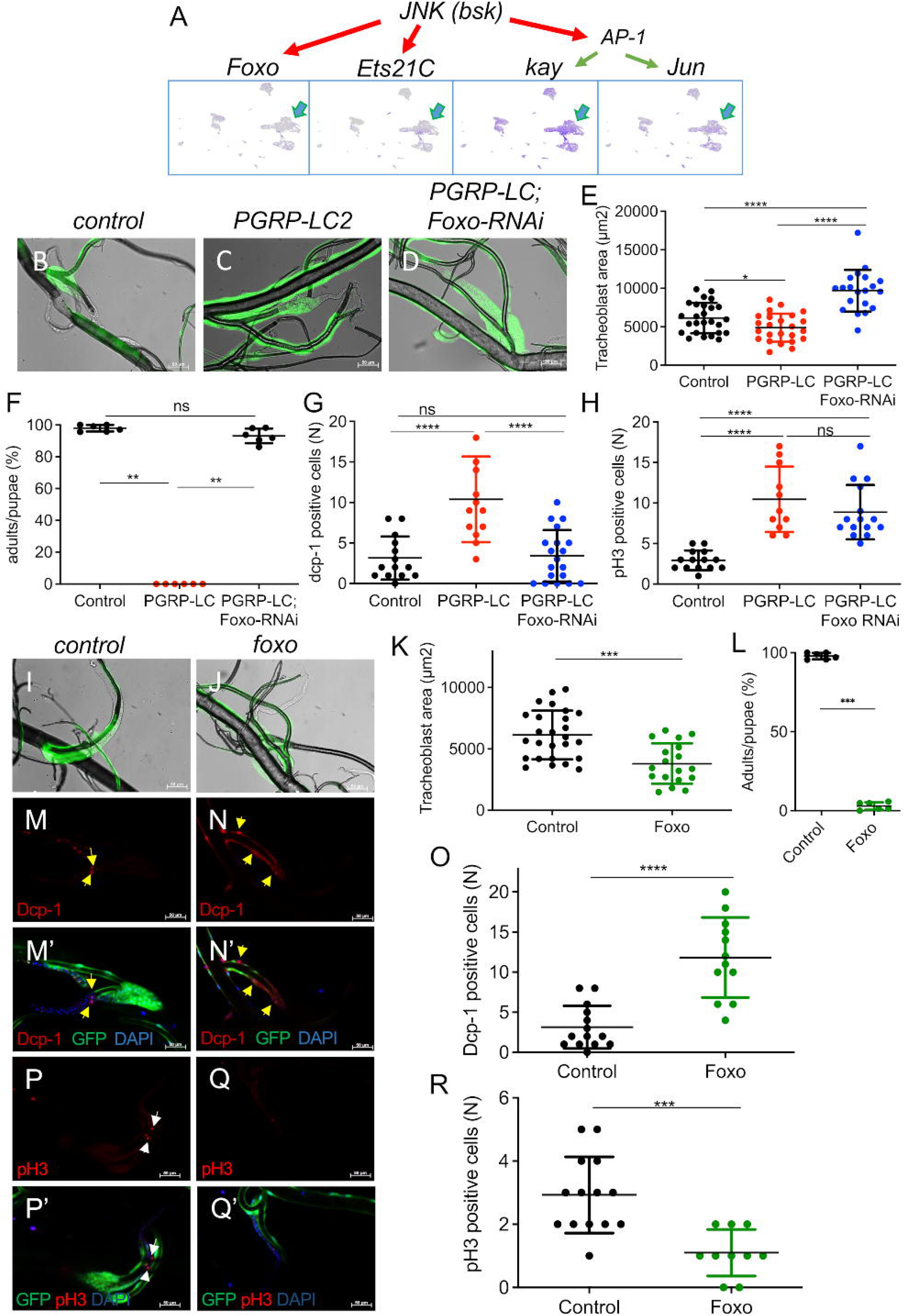
Foxo acts downstream of JNK and is necessary and sufficient for mediating the apoptotic signals in tracheoblasts. Dissected tracheae from control larvae (A, H, L, L’, O, O’; *btl-Gal4, UAS-GFP; tub-Gal80 [ts] > w^1118^*), of PGRP-LC overexpressing larvae (B; *Btl-Gal4, UAS-GFP; tub-Gal80 [ts] > UAS-PGRP-LC(2)*), those experiencing concurrent expression of PGRP-LC and foxo-RNAi (C; *Btl-Gal4, UAS-GFP; tub-Gal80 [ts] > UAS-PGRP-LC(2); UAS-foxo-RNAi)* and those overexpression of *foxo* (I, M, M’, P, P’; *btl-Gal4, UAS-GFP; tub-Gal80 [ts] > UAS-Foxo*). Adults to pupae ratios of animals shifted after 6d of development to the permissive temperature (29 °C), experiencing *PGRP-LC* overexpression or concurrent *PGRP-LC* and *Foxo*-RNAi overexpression (E), and of those experiencing *Foxo* overexpression (K). Anti-dcp1 signals (L, L’, M, M’) and anti-pH3 (O, O’, P, P’) signals are shown in red, GFP in green, and DAPI in blue. Yellow arrows (pH3 positive cells, white arrows) highlight positive cells. Statistical analysis of tracheoblast areas (D, J), of dcp1-signals (F, N), and pH3 signals (G, Q). Scale bar = 50 µm. Statistical significance was evaluated by Mann–Whitney test, ** p < 0.01, *** p < 0.001, ****p < 0.0001, n≥10.

### Signaling Downstream of JNK in Apoptosis and Proliferation

To dissect transcriptional mechanisms downstream of JNK, we focused on the three major canonical effectors of this pathway: Foxo, Ets21C, and the AP-1 components Kay (*Drosophila* Fos) and Jra (*Drosophila* Jun). All four factors were expressed in tracheoblasts (Fig. 5A), with single-cell RNA data confirming strong enrichment in the progenitor cell area (Fig. 5A, arrows) [35].

Given our previous finding that Foxo acts downstream of the IMD/JNK axis to regulate epithelial structure in the dorsal trunk [31], we examined its role in SB tracheoblast remodeling. Knockdown of *foxo* via RNAi completely rescued the PGRP-LC–induced reduction in tracheoblast area size (Fig. 5B–E). As expected, PGRP-LC overexpression alone markedly reduced SB tracheoblast size (Fig. 5B, C), while concurrent *foxo*-RNAi expression restored size to control levels (Fig. 5D, E).

We next tested whether *foxo* knockdown could also rescue the developmental lethality caused by PGRP-LC overexpression. Strikingly, simultaneous expression of *foxo*-RNAi and PGRP-LC fully restored adult viability (Fig. 5F).

Finally, we asked whether apoptosis and proliferation phenotypes triggered by IMD/JNK activation are Foxo-dependent. Dcp-1 staining revealed that *foxo*-RNAi co-expression completely abolished the PGRP-LC–induced increase in apoptotic SB tracheoblasts, reducing counts to control levels (Fig. 5G). In contrast, *foxo* knockdown failed to rescue the elevated rate of proliferation, as pH3-positive cell numbers were similar between PGRP-LC–only and PGRP-LC + *foxo*-RNAi animals (Fig. 5H).

In summary, Foxo acts downstream of the Tak1/JNK pathway to mediate apoptosis but not proliferation in SB tracheoblasts following IMD pathway activation.

### Foxo Overexpression in the Airway Epithelium Induces SB Tracheoblast Reduction via Apoptosis

Overexpression of foxo in the airway epithelium induced a dorsal trunk epithelial thickening phenotype [31]. We next tested whether *foxo* activation is sufficient—in addition to being necessary—to reduce SB tracheoblast area. Ectopic expression of *foxo* in the tracheal system reduced SB tracheoblast area to less than half of control values (Fig. 5I–K).

To confirm tracheoblast specificity, we used the progenitor cell-specific driver *ci*-Gal4, which restricts expression to tracheoblasts. Targeted *foxo* overexpression under *ci*-Gal4 markedly decreased T4 and T5 SB tracheoblast areas compared with *ci*-Gal4; *w1118* controls (Supplementary Fig. 5A–J), closely phenocopying the effect of whole-trachea overexpression. As with PGRP-LC overexpression, early developmental activation of *foxo* in the airway epithelium was lethal (Fig. 5L).

We next examined the mechanism underlying tracheoblast size reduction upon *foxo* overexpression. Apoptosis, assessed by Dcp-1 staining, was significantly elevated in T5 SB tracheoblasts of *foxo*-overexpressing animals (11.83 ± 1.44) compared to controls (3.14 ± 0.71; *p* < 0.0001; Fig. 5M–O). In contrast, proliferation, quantified by pH3 staining, was significantly reduced in T4 and T5 SB tracheoblasts expressing ectopic *foxo* (Fig. 5P–R).

These findings mirror the RNAi rescue experiments, confirming that while Foxo is both necessary and sufficient to mediate the IMD/JNK-driven apoptotic response in SB tracheoblasts, it does not contribute to the proliferative response triggered by IMD pathway activation.

### Ets21C is Required for Larval Tracheal Development

In addition to Foxo, the JNK pathway regulates other transcription factors, including Ets21C, previously shown to maintain intestinal homeostasis in *Drosophila* intestinal stem cells and enterocytes. Given that Foxo activation triggers apoptosis in tracheoblasts, we hypothesized that Ets21C might mediate IMD/JNK-driven tracheoblast proliferation by regulating the cell cycle and promoting cell division.

To test this, we employed a method where Ets21C was silenced in the trachea using *btl*-Gal4-driven RNAi. This was done in conjunction with PGRP-LC overexpression to activate the IMD pathway. Compared to controls, tracheoblast areas were markedly reduced at both 1 day and 2 days of PGRP-LC activation (Fig. 6A–D, A′–D′). Tracheal epithelial thickness in double-manipulated animals was comparable to PGRP-LC-only flies (Supplementary Fig. 6), suggesting that Ets21C selectively affects tracheoblasts but not epithelial cells. pH3-positive cell counts were almost completely rescued by Ets21C knockdown, approaching control levels under IMD/JNK activation (Fig. 6E–H).

Silencing Ets21C in the absence of PGRP-LC overexpression also resulted in smaller SB tracheoblasts (Fig. 6I–K) and significantly fewer pH3-positive cells (Fig. 6L) compared to controls, indicating a role for Ets21C in basal tracheoblast proliferation. Developmental timing experiments using the temperature-inducible TARGET system showed that RNAi activation during the first 4 days of development led to pupal lethality. In contrast, activation at the embryonic stage caused death during early L3. Structural analysis of these animals showed severe defects in tracheal architecture (Supplementary Fig. 7).

Conversely, overexpressing Ets21C in the trachea significantly increased tracheoblast size (Fig. 6M–O) and elevated pH3-positive cell counts (Fig. 6P). Together, these findings demonstrate that Ets21C is both necessary and sufficient for tracheoblast proliferation and that it plays an essential role in larval tracheal development.

### The transcription factor AP-1 is activated in the airway epithelium by IMD signaling, dependent on the JNK signaling pathway

To further clarify the signaling events downstream of the IMD/JNK axis, we examined AP-1, a canonical transcription factor that mediates the apoptosis-inducing effects of JNK pathway activation [36]. First, we aimed to elucidate whether AP-1 is activated via the IMD/JNK axis or by stressors known to activate the JNK pathway (Figure 7). Using the AP-1 reporter TRE-RFP [37], we found that activation of the IMD pathway substantially increased TRE-RFP signal (Figure 7A, B, D). To demonstrate that AP-1 signaling caused by IMD activation is JNK-dependent, we analyzed flies with the TRE-RFP reporter along with UAS-*PGRP-LC* and *Bsk^DN^* concurrently expressed in the trachea; this resulted in an almost complete rescue of the AP-1 signaling seen in the matching controls (Figure 7A, C, D). In addition, we explored whether AP-1 is also activated by external stressors believed to be mediated via JNK signaling. Here, cigarette smoke (Figure 7F, F’), cold exposure (Figure 7G, G’), and hypoxia (Figure 7H, H’) stimuli increased the TRE-RFP signal more than 15-fold compared to controls (Figure 7E, E’) without any stimulus (Figure 7I).

**Figure 6.**
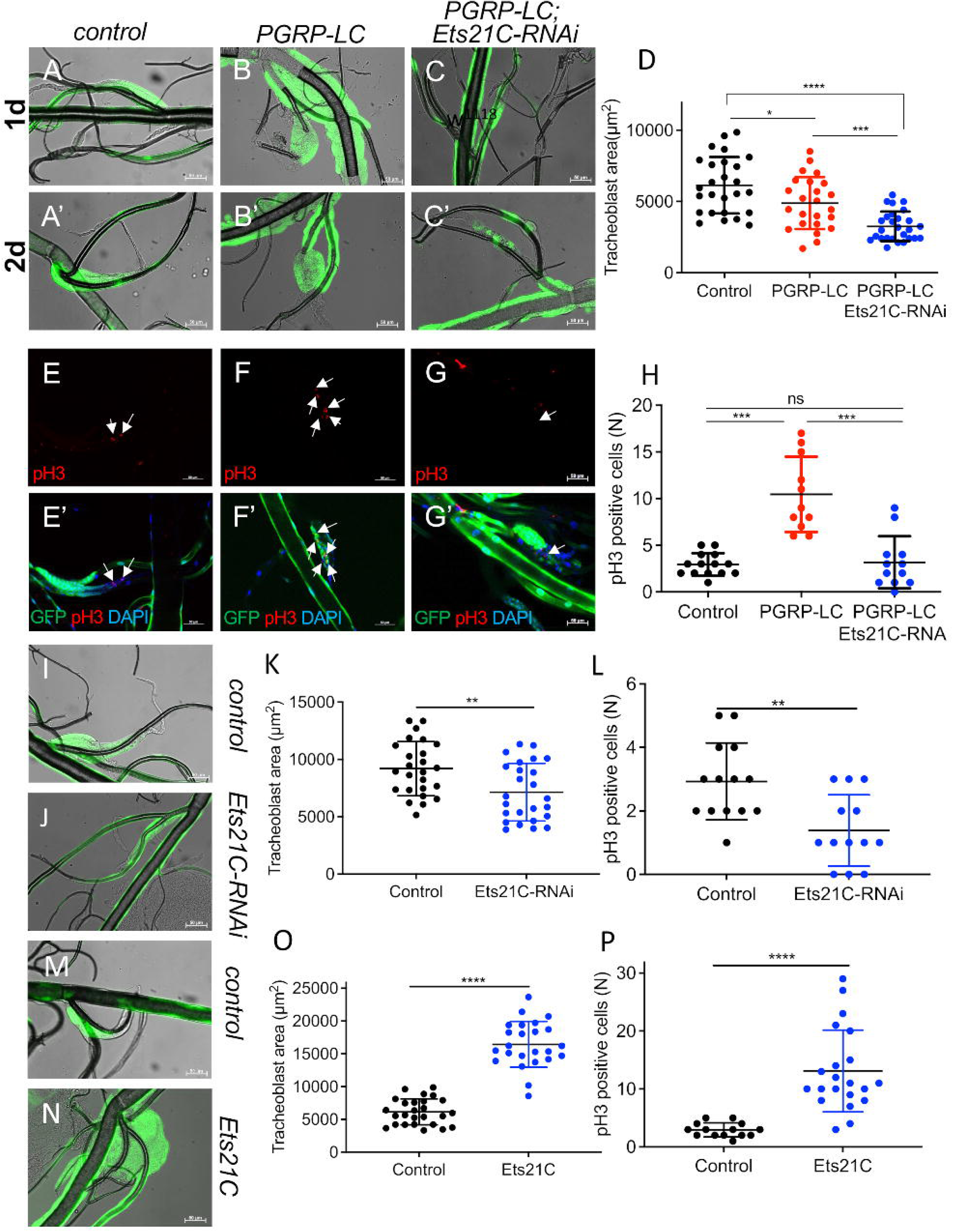
Ets21C enhances the adverse effects of PGRP-LC overexpression on tracheoblast size and is required for tracheoblast proliferation. (A-H) Dissected larvae of the control (A. A’, E, E’; *btl-Gal4, UAS-GFP; tub-Gal80[ts](btl.ts) > w^1118^*), those experiencing PGRP-LC overexpression (B, B’, F, F’; *btl-Gal4, UAS-GFP; tub-Gal80[ts](btl.ts) > UAS-PGRP-LC)* and those concurrent expression of *PGRP-LC* and *Ets21C*-RNAi (C, C’, G, G’*; btl-Gal4, UAS-GFP; tub- Gal80[ts](btl.ts) > UAS-PGRP-LC, UAS-Ets21C-RNAi).* Analysis of tracheoblast sizes (A-D) with quantitative analyses of tracheoblast sizes (D). Trachea stained with anti-PH3 (red)(E-H) with the quantitative evaluation of the numbers of PH3 positive cells (H). Embryos of *btl-Gal4, UAS-GFP; tub-Gal80[ts](btl.ts) > w^1118^*(black) and *btl-Gal4, UAS-GFP; tub-Gal80[ts](btl.ts) > UAS-Ets21C-RNAi* (green) were placed at 18 °C and transferred to 29 °C on different days of development. The development of pupae and adults. (I-P) Dissected larval tracheae were isolated from controls (I, M, M’; *btl-Gal4, UAS-GFP; tub-Gal80 [ts] > w^1118^*) from animals experiencing Ets21C-RNAi (J, N, N’; *btl-Gal4, UAS-GFP; tub-Gal80 [ts] > UAS-Ets21C RNAi*), and *Ets21C* overexpression ( (K, O. O’; *btl-Gal4, UAS-GFP; tub-Gal80 [ts] > UAS-Ets21C*). (L) Quantitative analysis of the area of T5 tracheoblasts. P, Quantitative analyses of the numbers of PH3 positive cells. Statistical significance was evaluated by Mann– Whitney test, *p < 0.05, **p < 0.01, ***p < 0.001. n≥10.

**Figure 7.**
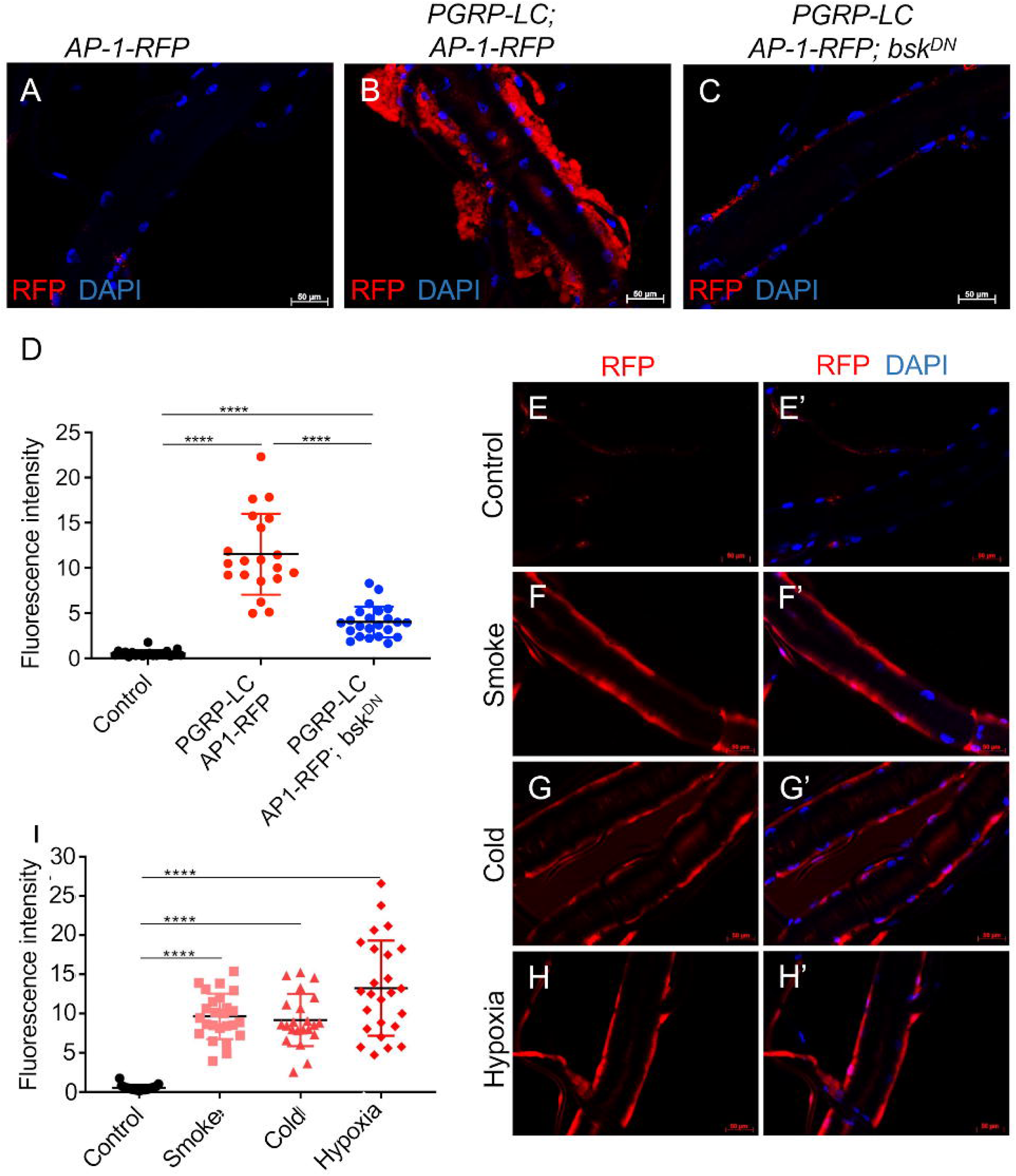
The JNK branch of the IMD signaling pathway mediates AP-1 activation in airway epithelial cells. (A) Fluorescence images of third-instar larvae expressing RFP-based AP-1 reporter and (B), expressing RFP-based AP-1 reporter and *PGRP-LC* simultaneously. (C) Expression of RFP-based AP-1 reporter, PGRP-LC, and Bsk^DN^ simultaneously. (E-H, E’-H’) Dissected trachea of late third instar larvae from AP1-RFP. (E, E’) without any treatment or stimulation as a control group. (F, F’) exposed to 2 cigarettes for 45 min. (G, G’) stored on ice for 12 h. (H, H’) exposed to 1% oxygen for 12 h. Tracheal tissue was simultaneously monitored for RFP (A-C, E-H, E’-H’); Nucleic DAPI staining is shown in blue (A-C, É-H’). (D, I) Fluorescence intensity of AP1 was quantified by Image J. Scale bar = 50 µm. Statistical significance was evaluated by Mann–Whitney test, ****p < 0.0001. n≥20.

### Blocking Kay or Jra rescued the PGRP-LC overexpression-induced reduction in tracheoblast area, but did not affect the tracheal epithelial thickening phenotype

Next, we evaluated whether AP-1 is necessary for the IMD-activation-induced phenotype of smaller tracheoblast areas. We blocked AP-1 formation by reducing one of the two AP-1 constituents through *Kay^DN^* overexpression (Figure 8A–D). This was conducted concurrently with IMD-pathway activation via ectopic *PGRP-LC* expression (Figure 8C, D). We observed a rescue of the *PGRP-LC*-induced tracheoblast phenotype (Figure 8A–D), but not the PGRP-LC-induced tracheal epithelial thickening (Supplementary Figure 8I–K).

**Figure 8.**
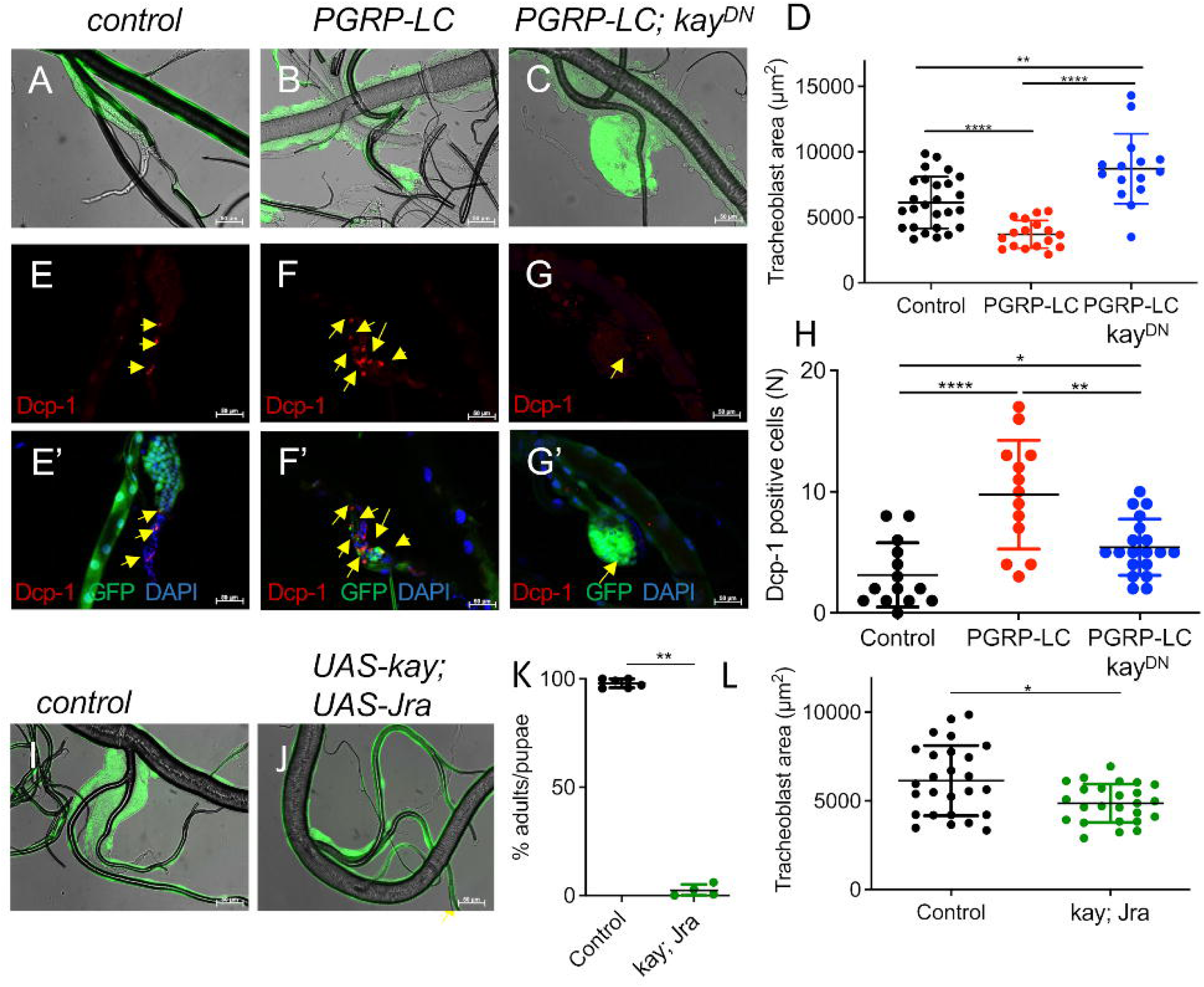
Blocking AP-1-signaling rescues PGRP-LC-induced reduced size of tracheoblasts. Dissected tracheae from *btl-Gal4, UAS-GFP; tub-Gal80 [ts] > w^1118^* control larvae (A, E, E’, I). *Btl-Gal4, UAS-GFP; tub-Gal80 [ts] > UAS-PGRP-LC1* larvae with ectopic expression of PGRP-LC in the trachea (B, F, F’). *Btl-Gal4, UAS-GFP; tub-Gal80 [ts] > UAS-PGRP-LC1; UAS-Kay^DN^* larvae with concurrent ectopic expression of PGRP-LC and Kay^DN^ in the trachea (C, G, G’). (A-D) T5 tracheoblast areas. (D) Quantitative analysis of concurrent expression of Kay^DN^ and PGRP-LC in the trachea of T5 tracheoblast areas. Dissected tracheae from control larvae (E, E’,; *btl-Gal4, UAS-GFP; tub-Gal80 [ts] > w^1118^*), PGRP-LC1 larvae (F, F’; *btl-Gal4, UAS-GFP; tub-Gal80 [ts] > UAS-PGRP-LC1*) and PGRP-LC1; Kay^DN^ larvae (G, G’; *btl-Gal4, UAS-GFP; tub-Gal80 [ts] > UAS-PGRP-LC1; UAS-Kay^DN^*) were stained with a Dcp-1 antibody and the signal is shown in red. Tissue of trachea is stained GFP in green, nuclei are stained DAPI in blue (É-G’). Yellow arrows highlight Dcp-1 positive cells. (H) Statistical analysis of the dcp-1 positive cells of T5 tracheoblasts. Effects of overexpression of AP-1 (UAS-Kay; UAS-Jra) (J) and matching controls (I). (K) Adult to pupae ratio of control animals and those experiencing AP-1 overexpression. (L) Statistical analysis of tracheoblast areas. Scale bar = 50 µm. Statistical significance was evaluated by Mann–Whitney test, ns, not significant, *p < 0.05, **p < 0.01, ***p < 0.001, ****p < 0.0001. n≥10.

To understand the underlying mechanism, we calculated the number of apoptotic cells and found that co-expressing *Kay^DN^* along with *PGRP-LC* rescued tracheoblasts from apoptosis to control levels (Figure 8E–H). Here, the level of Dcp-1-positive cells was only slightly higher than in controls, yet it was significantly reduced compared with *PGRP-LC* overexpression alone (Figure 8H). Seeking to determine whether AP-1 is not only necessary but also sufficient to induce the phenotype seen in IMD-pathway-activated tracheoblasts, we first overexpressed Kay or Jra individually in the tracheal system (Supplementary Figure 8). We found that in T4 and T5 tracheoblasts, Kay or Jra overexpression resulted in a slight increase in size rather than the expected decrease. Therefore, Jra and Kay were ectopically overexpressed together in the airways to allow functional AP-1 activation (Figure 8I, J). As with *Drosophila* overexpressing *PGRP-LC* (Figure 2A) or *foxo*, activation of AP-1 after 6 days of development in the tracheal system also led to the death of the animals (Figure 8K). Overexpression of this functional AP-1 resulted in smaller tracheoblast areas, similar to the effects observed following IMD pathway activation (Figure 8I, J, L).

To analyze the exact positioning of Foxo and AP-1 in the signal chain downstream of JNK, we introduced the TRE-RFP reporter into flies overexpressing *foxo* in the trachea to determine whether *foxo* could activate AP-1. RFP was markedly increased in the tracheal epithelial cells of *Drosophila* overexpressing *foxo* (1.48 ± 0.17) compared to the control group (0.93 ± 0.19, p < 0.0001) (Figure 9A–C). Although the increase was slight, it was statistically significant. This indicates that ectopic overexpression of *foxo* in airway epithelial cells induces activation of AP-1. To address the question of whether the positioning of AP-1 downstream of *foxo* is also seen in functional assays, we generated *Drosophila* flies in which *foxo* overexpression was targeted to the trachea concurrently with silencing of *Jra* (*Jra-RNAi*) to inhibit the generation of AP-1 (Figure 9D–H). Overexpression of *foxo* in the airway system early in animal development leads to animal death. After 6 days of incubation at 18 °C, 3.0 ± 0.97% of animals overexpressing *foxo* developed into adults, while 48.76 ± 2.27% of animals overexpressing *foxo* concurrently with Jra-RNAi developed into adults (Figure 9D). As shown in Figure 9E–K, the reduction in tracheoblast areas caused by *foxo* overexpression was fully rescued by concurrent expression of *Jra-RNAi*, resulting in slightly larger areas than those in untreated controls. These results (Figure 9) suggest that the AP-1 transcription factor is both sufficient and necessary for *foxo*-induced apoptosis.

**Figure 9.**
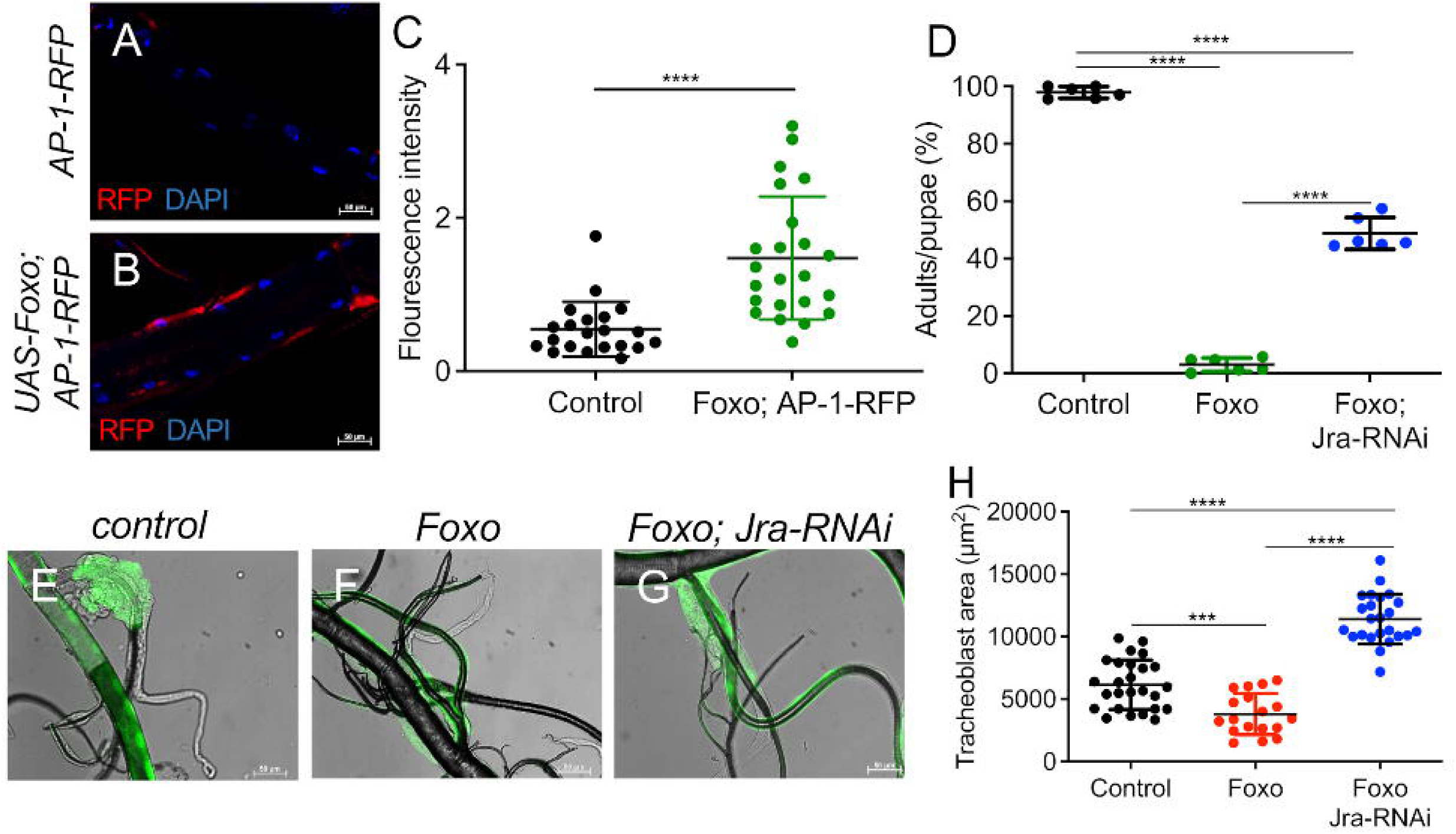
Foxo-induced apoptosis requires the AP1 transcription factor. (A) Fluorescence images of third-instar larvae expressing RFP-tagged AP1 (*ppk4-Gal4 x AP1-RFP*). (B) Overexpressing RFP-tagged AP1 and FoxO simultaneously (*ppk4-Gal4 x UAS-FoxO; AP1-RFP*). Tissue of trachea analysed for RFP in red, nuclei (DAPI) in blue (A, B). (C) A quantitative analysis of the fluorescence intensity of AP1 was performed with Image J. Scale bar = 50 µm. Statistical significance was evaluated by Mann–Whitney test, ****p < 0.0001. n ≥ 20. (D) Statistical analysis of the adult rate of larvae incubated at 18 °C for 6 days and then transferred to 29 °C for continued incubation. Statistical significance was evaluated by Mann–Whitney test, ns, not significant, **p < 0.01, n = 5. (E) Dissected tracheae from *btl-Gal4, UAS-GFP; tub-Gal80 [ts] > w^1118^* control larvae. (F) Dissected tracheae from *btl-Gal4, UAS-GFP; tub-Gal80 [ts] > UAS-Foxo*. (G) Dissected tracheae from *btl-Gal4, UAS-GFP; tub-Gal80 [ts] > UAS-Foxo; UAS-Jra-RNAi*. (H) Quantitative analysis of the area of T5 tracheoblasts. Scale bar = 50 µm. Statistical significance was evaluated by Mann–Whitney test, *p < 0.05, **p < 0.01, ***p < 0.001, ****p < 0.0001. n≥10.

The investigations carried out here have clarified the signalling pathway linking immune activation to apoptosis and proliferation in the precursor cells of the respiratory organs (Figure 10). The crucial link between immune signalling and the stress-sensitive JNK signalling pathway, established by Tak1, is a key finding. Notably, the signals for proliferation and apoptosis below JNK diverge into distinct pathways. The interaction between Foxo and AP-1 in apoptosis induction, however, remains a mystery.

**Figure 10:**
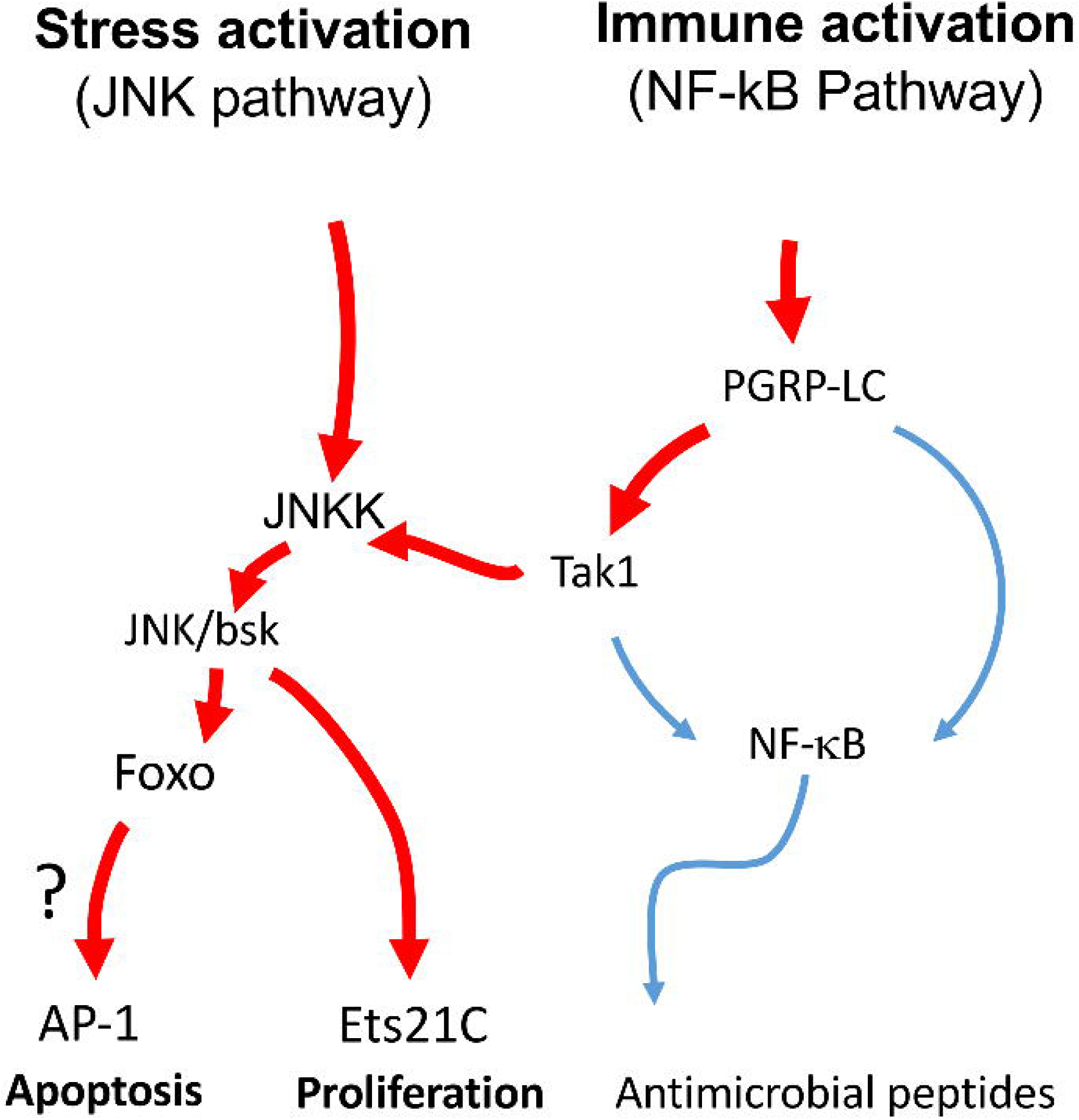
Scheme of the signaling transduction pathways mediating the effects of apoptosis and proliferation in response to immune activation. Immune (IMD-signaling) and Stress-pathway activation induce both apoptosis and proliferation by targeting JNK signaling.

## Discussion

### Chronic Inflammation Reshapes Progenitor Cell Fate

Inflammation is indispensable for clearing infections and maintaining tissue integrity. However, when sustained, it becomes detrimental—disrupting progenitor function, depleting regenerative niches, and driving disease pathology [38]. Our study demonstrates that persistent immune activation in the *Drosophila* airway elicits a dual response in tracheal progenitor cells: apoptosis and compensatory proliferation. These changes occur through cell-intrinsic mechanisms, not as secondary effects from immune effector cells. The result is niche shrinkage, compromised regenerative capacity, and altered epithelial homeostasis.

Such duality—where immune stress triggers both proliferation and cell death—is also observed in other systems [13, 39–41], including intestinal stem cells and mammalian lung progenitors, reinforcing the idea that chronic inflammation and regeneration are intertwined through shared signaling logic.

### Tracheoblasts Are Immune-Competent Progenitors

We establish that *Drosophila* tracheal progenitors (tracheoblasts) are not passive recipients of immune signals; instead, they are immune-competent, capable of recognizing microbial patterns and producing antimicrobial peptides. This property mirrors intestinal stem cells in *Drosophila* [42, 43] and basal or AT2 cells in the mammalian lung [44, 45], which also exhibit direct immune engagement.

This immune competence correlates with niche architecture. In the intestine, mammalian stem cells are protected by Paneth cells that produce AMPs and mediate host-microbe interactions [46, 47]. In contrast, airway progenitors—like tracheoblasts, basal cells, and AT2 cells—exist in unshielded environments, requiring them to defend themselves. This makes them vulnerable to chronic immune activation, which over time can deteriorate their regenerative function and lead to disease, such as COPD [21, 22].

### JNK Pathway: A Central Hub Integrating Stress and Immune Signals

Our data reveal a signaling bifurcation in response to chronic activation of the IMD pathway. Rather than acting via canonical NF-κB (Relish) signaling, the immune input is redirected at Tak1 to activate the JNK pathway [33], which orchestrates fate decisions in progenitor cells. This redirection enables JNK to serve as an integrator of immune and stress signals—a theme consistent with its role in regulating stemness, differentiation, and apoptosis across tissues [48–50].

Importantly, JNK activation alone was sufficient to induce both apoptosis and proliferation, while blocking JNK rescued immune-induced tissue shrinkage, confirming its centrality. Moreover, JNK acts as a real signal integrator, as targeting this pathway through canonical TNF-activation induced exactly the same phenotypes as observed in response to IMD-pathway activation.

### Ets21C Mediates Proliferation via JNK in Immune-Stressed Progenitors

We identify Ets21C, a conserved ETS transcription factor, as a key effector of immune-induced proliferation downstream of JNK. This aligns with prior findings where Ets21C is induced by infection, injury, and oncogenic stimuli [51–55], and plays roles in intestinal stem cell renewal and epithelial tumorigenesis.

Ets21C is functionally analogous to human FLI1 and ERG, both of which regulate lung development and fibrosis [56–60]. Our data show that Ets21C is essential not only for tracheoblast proliferation during normal development but also for compensatory proliferation under immune challenge. These findings establish Ets21C as a context-dependent proliferative switch—a potential target for modulating regeneration in inflammatory disease [61, 62].

### Foxo and AP-1 Cooperate to Drive Apoptosis Under Immune Stress

Alongside Ets21C-mediated proliferation, Foxo and AP-1 (Jra/Kay) mediate the apoptotic arm of the response. Both are necessary and sufficient for inducing apoptosis in airway progenitors during chronic inflammation. Foxo, a known integrator of metabolic, stress, and immune signals [63–68], is central to maintaining stem cell quiescence in both Drosophila and mammals [69, 70]. However, under sustained immune stress, Foxo shifts from a guardian of quiescence to a driver of cell death, depleting the progenitor pool.

AP-1, a canonical JNK target, is similarly involved in apoptosis and stem cell regulation [71, 72]. Our results suggest that Foxo and AP-1 may act sequentially or synergistically downstream of JNK, mediating apoptosis specifically in progenitor cells, but not in mature epithelial cells—where AP-1 is dispensable.

### Modular Signaling Architecture Enables Independent Tuning of Fate

One of the most striking findings is the modularity of the immune-induced fate program. Blocking Foxo or AP-1 rescues apoptosis but not proliferation, while knocking down Ets21C prevents proliferation but not cell death. This functional separation allows for the independent modulation of each arm of the stress response, enabling the tissue to balance regeneration and elimination during immune challenge.

This architecture has broad implications. In pathologies like pulmonary fibrosis or chronic bronchitis, similar signaling logics may underlie epithelial turnover, progenitor exhaustion, and fibrosis. Targeting individual arms of the response, e.g., selective inhibition of Ets21C or AP-1, could allow us to modulate regeneration without impairing defense.

### Clinical and Therapeutic Implications

Ets21C’s human homologs—FLI1 and ERG—are already implicated in vascular remodeling, fibrosis, and lung cancer [56–60], and are considered druggable transcription factors [61, 62]. Our findings provide mechanistic insight into how these factors function during inflammation-induced regeneration, offering a basis for future therapeutic exploration in chronic lung diseases. Similarly, manipulating Foxo or AP-1 [73–75] may allow for the preservation of progenitor identity under immune stress, preventing niche exhaustion and organ failure.

## Conclusion

Chronic inflammation imposes a fundamental challenge to tissue homeostasis by coupling defense mechanisms to stress-induced remodeling. Here, we define a conserved signaling logic whereby immune activation converges on JNK to bifurcate progenitor fate into apoptotic and proliferative branches via Foxo/AP-1 and Ets21C, respectively. This modular organization enables epithelial progenitors to tune their regenerative capacity while limiting excessive growth under persistent immune stress. Notably, human orthologs of these transcriptional regulators—FOXO3A, JUN/FOS, and ERG/FLI1—are key mediators of lung regeneration, fibrosis, and inflammation, implying evolutionary conservation of this regulatory circuit. We propose that chronic activation of this axis underlies progenitor depletion and maladaptive remodeling seen in airway diseases such as COPD and pulmonary fibrosis. By delineating how JNK integrates immune and stress inputs to dictate progenitor outcomes, this study provides a blueprint for targeted strategies to restore regenerative balance in chronically inflamed epithelia.

## Materials and Methods

| Primary antibodies: | Vendor |
| --- | --- |
| α-phospho-Histone H3 (Ser10), rabbit | Cell Signaling Technology (Danvers, USA)<br>Cat # 53348T |
| α-beta-galactosidase (40-1a), mouse | Developmental Studies Hybridoma Bank (DSHB),<br>(Iowa City, USA), RRID: AB_528100 |
| α-Dcp1, rabbit | Cell Signaling Technology (Danvers, USA)<br>Cat # 9578S |
| α-GFP, mouse (12A6-b) | DSHB (Iowa City, USA) |
|  | RRID: AB_2617417 |
| **Secondary antibodies:** |  |
| α-mouse-AlexaFluor 488, goat | Jackson ImmunoResearch Laboratories (West Grove, USA) |
| α-rabbit-AlexaFluor 488, goat | Life Technologies (Carlsbad, USA) |
| α-mouse-AlexaFluor 555, goat | Cell Signaling Technology (Danvers, USA) |
| α-rabbit-Cy3 | Jackson ImmunoResearch Laboratories (West Grove, USA) |
| **Fluorescent dyes/ conjugated proteins** |  |
| Roti®-Mount FluorCare DAPI (4,6-diamidino-2-phenylindole) | Carl Roth (Karlsruhe, Germany) |

| **Experimental models: Organisms/strains** | **Vendor** |
| --- | --- |
| *Drosophila melanogaster* w<sup>1118</sup> | BDSC (3605) |
| *Drosophila melanogaster* UAS-PGRP-LE | BDSC (33054) |
| *Drosophila melanogaster* UAS-PGRP-LCx | BDSC (30919) |
| *Drosophila melanogaster* UAS-PGRP-LCx | BDSC (30918) |
| *Drosophila melanogaster* UAS-FoxO II | BDSC (9575) |
| *Drosophila melanogaster* UAS-FOXO-RNAi | BDSC (32993) |
| *Drosophila melanogaster* UAS-FOXO-RNAi | BDSC (27656) |
| *Drosophila melanogaster* UAS-Kay-DN | BDSC (7214) |
| *Drosophila melanogaster* UAS-Jra-RNAi | BDSC (31595) |
| *Drosophila melanogaster* UAS-Jra | BDSC (7216) |
| *Drosophila melanogaster* UAS-fra (Kay) | BDSC (7213) |
| *Drosophila melanogaster* Jra-GFP | BDSC (50755) |
| *Drosophila melanogaster* Kay-GFP | BDSC (38657) |
| *Drosophila melanogaster* P(UAS-bsk.DN) [76]2 | BDSC (6409) |
| *Drosophila melanogaster* UAS-Rel-V5 | BDSC (55779) |
| *Drosophila melanogaster* UAS-Rel-RNAi | BDSC (28943) |
| *Drosophila melanogaster* UAS-Ets21c-RNAi | BDSC (39069) |
| *Drosophila melanogaster* UAS-RelD | S. Cherry, Univ. Pennsylvania, USA [77] |
| *Drosophila melanogaster* dTAK1DN;PGRP-LCx | generated by ourselves [34] |
| *Drosophila melanogaster* Ci-gal4, UAS-GFP; gal80ts | Xavier Franch-Marro, Barcelona, Spain |
| *Drosophila melanogaster* btl-Gal4, UAS-GFP; | Leptin Group Heidelberg |
| Gal80ts |  |
| *Drosophila melanogaster* Balancer (2<sup>nd</sup> + 3<sup>rd</sup>) | T. Stork, Univ. Massachusetts, Worcester, USA |
| *Drosophila melanogaster* TRE (AP1)-RFP attP40 | [78] |
| *Drosophila melanogaster* ppk4-gal4 | M. Welsh, Iowa, USA [79] |
| *Drosophila melanogaster* UAS-basketDN; PGRP-LCx | generated by ourselves [34] |
| *Drosophila melanogaster* UAS-PGRP-LCx; UAS-Foxo-RNAi | generated by ourselves [34] |
| *Drosophila melanogaster* Sp(BL)/CyO; TM2/TM6 | T. Stork, Univ. Massachusetts, Worcester, USA |
| *Drosophila melanogaster* UAS-Ets21c |  |

### Experimental models: *Drosophila*

The following transgenic *Drosophila* strains used were supplied by the Bloomington *Drosophila* Stock Center (BDSC): w^1118^ (BDSC 3605), UAS-PGRP-LC1 (BDSC 30919), UAS-PGRP-LC2 (BDSC 30918), UAS-PGRP-LE (BDSC 33054), UAS-basketDN (BDSC 6409), UAS-foxo1 (BDSC 9575), UAS-foxo2 (BDSC 43633), UAS-foxoRNAi1 (BDSC 32993), foxoRNAi2 (BDSC 27656), UAS-kayDN (BDSC 7214), UAS-fra (Kay) (BDSC 7214), Kay-GFP (BDSC 38657), UAS-Jra (BDSC 7216), UAS-JraRNAi (BDSC 31595), Jra-GFP (BDSC 50755), UAS-Rel-V5 (BDSC 55779), UAS-Rel-RNAi (BDSC 28943).

In addition, the following stocks used were generously provided by scientists in the *Drosophila* research community: UAS-relD (S. Cherry, Univ. Pennsylvania, USA; [77], btl-Gal4, tubP-Gal80ts, UAS-GFP (M. Leptin, Heidelberg, Germany), Ci-gal4, UAS-GFP; gal80ts (Xavier Franch-Marro, Barcelona, Spain), ppk4-gal4 (M. Welsh, Iowa, USA [79], TRE (AP1)-RFP attP40 [78]. If not otherwise noted, stocks and experimental flies were raised on standard cornmeal-agar medium at 25 °C at a relative humidity of at least 60% with a 12 h:12 h light/dark cycle.

### Crossings

We used the binary Gal4/UAS expression system [80] to induce gene expression of interest in larval airways. To generate traditional Gal4/UAS and TARGET crosses, virgin females of the Gal4 line were collected and, at 3 to 5 days of age, crossed with 3- to 7-day-old males of the UAS line at 25 °C. Experiments were performed with late L3 larvae of the F1 generation; larvae were not sorted by sex. To activate the TARGET system, larvae were maintained at 18 °C (restrictive temperature) until reaching the desired developmental stage, then shifted to 29 °C (permissive temperature) for 2 days. All animal studies complied with the state animal protection rules of Schleswig-Holstein, Germany.

### Developmental Viability

For developmental viability analyses, eggs were collected for 24 hours and were not physically handled in any way. Each group had three replicates of more than 50 eggs. To ensure larval developmental viability, tub-Gal80[ts] was used to limit UAS responder expression at the larva stage. Animals were raised at 18 °C to keep the UAS responder gene silent. Animals at different days of development were transferred to 29 °C.

### Quantification of the epithelial thickness of T9 and measurement of the area of tracheoblasts

Later L3 larvae were washed in PBS to remove the remaining medium before drying on a paper towel. The tracheae of wandering larvae were dissected and the dorsal trunk of T9 and the tracheoblasts were selected for microscopy and quantification using Z-stack analysis. Z-stack images were documented in DIC and GFP channels with the Axio Imager.Z1 with Apo Tome (Zeiss). Measurements were performed using the AxioVision SE64 Rel. 4.9 Software (Zeiss). Statistical analyses were performed with GraphPad Prism7 (GraphPad Software).

### Infection of the larval tracheal system

Natural infection of larvae was performed as described [27, 81]. The gram-negative bacterium *Pectobacterium carotovorum* (Ecc-15, 2141) was cultured in LB broth overnight at 30°C. The bacteria were pelleted by centrifugation at 3200 × g for 20 min, resuspended in a small amount of LB and the absorbance measured at OD600. Next, 200 µl of the bacterial solution (OD600 = 120) was added to a vial containing developing 2nd and early 3rd instar larvae (3 days after egg laying) in standard cornmeal medium. After 24 h, the tracheae of 3rd instar larvae were used for microscopy.

### Preparation, fixation, and immunofluorescence staining

Late L3 larvae were washed first in PBS and then in 70% ethanol at room temperature before they were placed in a block dish containing PBS. Larvae were opened longitudinally; the airways were exposed, and all surrounding organs were removed. Tissues were fixed in 4% paraformaldehyde (dissolved in PBS) at room temperature for 45 minutes, followed by washing in PBS. For immunostaining, samples were permeabilized with PBS/0.1% Triton X-100 and incubated with PBS/0.1% Triton X-100/10% normal goat serum (Sigma-Aldrich) for 60 minutes to block nonspecific binding sites. The tissue was then incubated overnight with primary antibody solved in PBS/0.1% Triton X-100/10% NGS at 4 °C. GFP signals were amplified by immunostaining with polyclonal rabbit anti-GFP (1:500, Sigma-Aldrich, Merck KGaA, Darmstadt, Germany). A monoclonal rabbit Cleaved *Drosophila* Dcp1 (1:200, Cell Signaling, Frankfurt/M, Germany) was used to detect apoptotic cells. Anti–phospho-histone H3 (Ser10; Cell Signaling Technology, 1:100) antibodies were used to detect mitotic cells. For the identification of beta-galactosidase in the larval airway system, monoclonal mouse anti-beta-galactosidase-antibody (40-1a) (Developmental Studies Hybridoma Bank, Iowa, USA) was used in a dilution of 1:150.

After primary antibody incubation, tissue was washed in PBS/0.1% Triton X-100 and incubated with secondary antibodies AlexaFluor 488-conjugated goat anti-mouse IgG or AlexaFluor 488-conjugated goat anti-rabbit IgG (Thermo Fisher Scientific/Invitrogen, Karlsruhe, Germany) in a dilution of 1:500 (dissolved in PBS/0.1% Triton X-100) at 4 °C overnight. Samples were washed twice in PBS/0.1% Triton X-100, then washed and stored in PBS. Nuclei were stained with 4’, 6-diamidino-2-2phenylindole, dihydrochloride (DAPI) (Roth, Karlsruhe, Germany, 6843). Specimens were analyzed and digital images were captured with a conventional fluorescence microscope (ZEISS Axio Imager Z1, Zeiss, Oberkochen, Germany).

### Cigarette smoke exposure

The vials containing animals were capped with a monitoring grid to allow the cigarette smoke to diffuse into the vial. Common research 1R6F cigarettes (CTRP, Kentucky University, Lexington, USA) were used for the experiments. For the smoke experiments, L3 larvae were exposed to two cigarettes smoked one time for 45 minutes. They were dissected after 8 hours of recovery.

### Cold and hypoxic exposure

In the experiment of larvae stimulated by cold, L3 larvae were placed in tubes containing food buried in ice for 12 hours. They were dissected after 8 hours of recovery. Larval hypoxia experiments were performed by exposing L3 larvae in tubes containing food to 1% oxygen in a desiccator for 12 hours. The trachea were dissected after 8 hours of recovery.

### Synchrotron radiation micro-computer tomography

Fly larvae have been imaged using synchrotron radiation-based x-ray micro-computed tomography (SRµCT). Imaging was performed at the Imaging Beamline P05 (IBL) [82–85] operated by the Helmholtz-Zentrum Hereon at the storage ring PETRA III (Deutsches Elektronen Synchrotron – DESY, Hamburg, Germany). Specimen were mounted by covering and attaching them to a toothpick using UV-hardening glue. Imaging was performed at an photon energy of 33 keV and an sample to detector distance of 100 mm. Projections were recorded using a custom developed 20 MP CMOS camera system [86] with an effective pixel size of 0.64 µm. For each tomographic scan 1501 projections at equal intervals between 0 and π have been recorded. Tomographic reconstruction has been done by applying a transport of intensity phase retrieval approach and using the filtered back projection algorithm (FBP) implemented in a custom reconstruction pipeline [87] using Matlab (Math-Works) and the Astra Toolbox [88, 89]. For processing, the raw projections were binned twice, resulting in an adequate pixel size of the reconstructed volume of 1.29 µm.

### Quantitative and statistical analysis

All experiments were repeated at least three times. Unpaired *t*-test followed by Holm multiple comparison testing, two-tailed Mann-Whitney *U* test, and one-way ANOVA were used to assess statistically significant differences. Statistical tests for individual experiments are shown in the figure legends, and the exact values of *n* are also indicated within the figures. Prism 7 (GraphPad) was used for graph editing and the statistical analyses; Olympus cellSens (version 1.16) and Leica Application Suite Advanced Fluorescence (version 2.7.3.9723) were used to take and edit photographs; and image processing was done using ImageJ (version 1.51j8).

## Acknowledgements

This work was supported by grants from the Deutsche Forschungsgemeinschaft (DFG; CRC1182, Project C2). We thank Britta Laubenstein and Christiane Sandberg for excellent technical assistance. Moreover, we thank Maria Leptin (Heidelberg, Germany), Tobias Stork (Oregon Health and Science University, USA), Xavier Franch-Marro (Barcelona, Spain), Mike Welsh (Iowa City, USA), and the Bloomington Stock Center at Indiana University (BDSC) for flies. We also acknowledge provision of beamtime related to the proposal BAG-20211055 at beamline P05, PETRA III at Deutsches Elektronen-Synchrotron (DESY), Hamburg, a member of the Helmholtz Association (HGF). This research was supported in part through the Maxwell computational resources operated at Deutsches Elektronen-Synchrotron DESY, Hamburg, Germany.

## Data availability

Data used for this publication are included in the manuscript (and supplementary information).

## Contributions

LS, XN, KZ, KS, JUH, and JB performed experiments and analyzed data. LS, HH, IB, SK-E, JB, and TR panned the project and drafted the manuscript. All authors read and corrected the final version of the manuscript.

## Ethics declaration Competing interests

The authors declare no competing interests.

