## Supplemental Figures 1-9 for "JNK integrates immune and stress signals to balance apoptosis and proliferation in airway progenitors"

Supplementary Figure 1

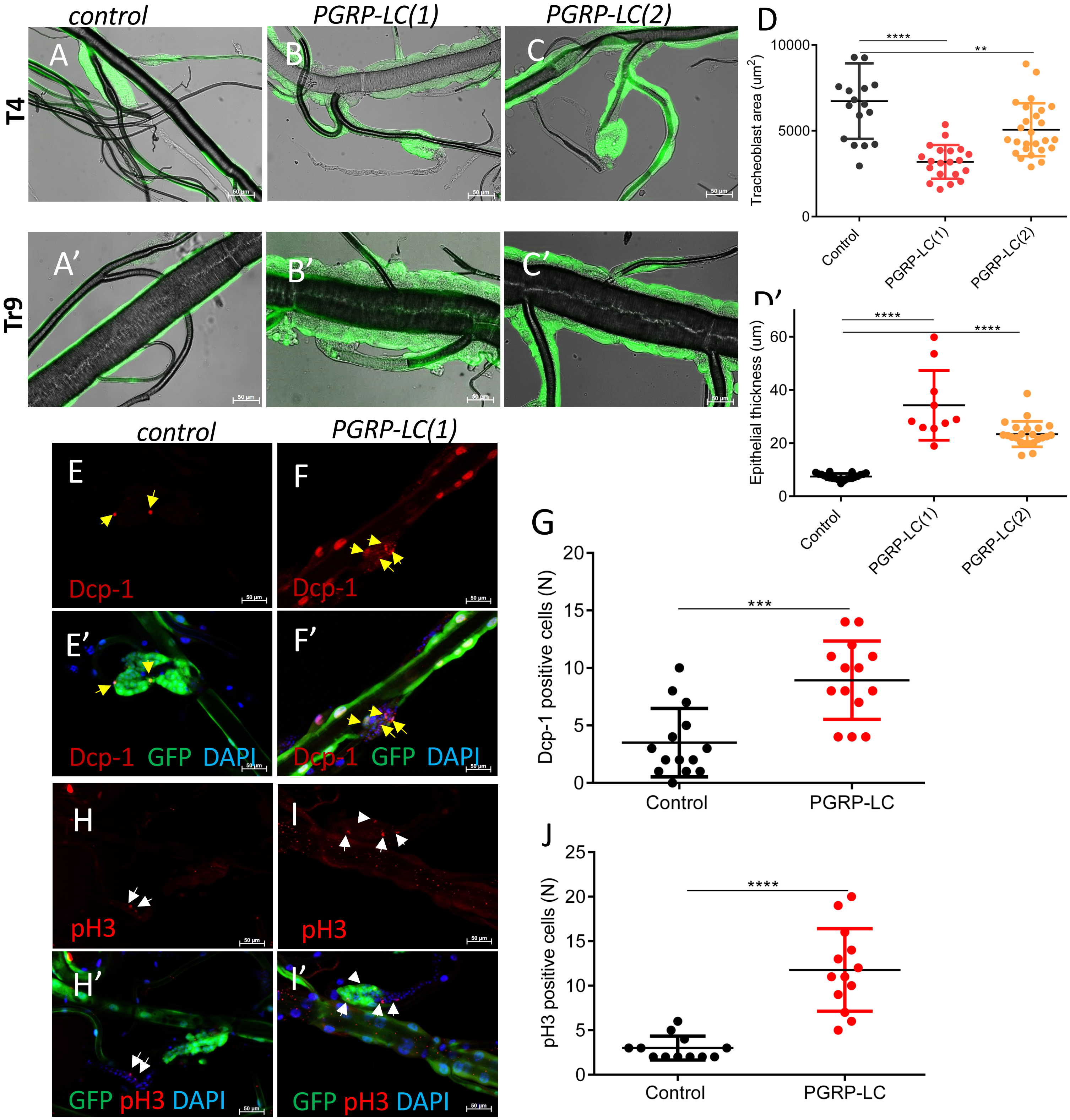


**Supplementary Figure 1. Constitutive activation of IMD signaling in the *Drosophila* airways induces considerable thickening of the airway epithelium and reduces the size of T4 SB tracheoblasts.** Dissected tracheae from control larvae (A, A', E, E', H, H'; *btl-Gal4, UAS-GFP; tub-Gal80 [ts] > $w^{1118}$*), PGRP-LC overexpressing larvae (B, B', F, F', I, I'; *btl-Gal4, UAS-GFP; tub-Gal80 [ts] > UAS-PGRP-LC(1),* and C, C'; *Btl-Gal4, UAS-GFP; tub-Gal80 [ts] > UAS-PGRP-LC(2)*). (D) Quantitative analysis of the area of T4 tracheoblasts; (D') Quantitative analysis of the thickness of the Tr9 airway epithelium. (E, E', F, F') Specimens were stained with the Dcp-1 antibody (red). (H, H', I, I') Samples were stained with the anti-pH3 antibody (red). (E', F', H'. I') The tissue of the trachea showing GFP is labeled green, and of the nuclear DAPI staining is blue. Yellow arrows highlight Dcp-1 positive cells. White arrows show pH3-positive cells. (G) Statistical analysis of the Dcp-1 positive cells of T4 tracheoblasts. (J) Statistical analysis of pH3 positive cells of T4 tracheoblasts. Scale bar = 50 μm. Statistical significance was evaluated by the Mann–Whitney test, $^{**}p < 0.01$, $^{***}p < 0.001$, $^{****}p < 0.0001$. n≥10.

### Supplementary Figure 2

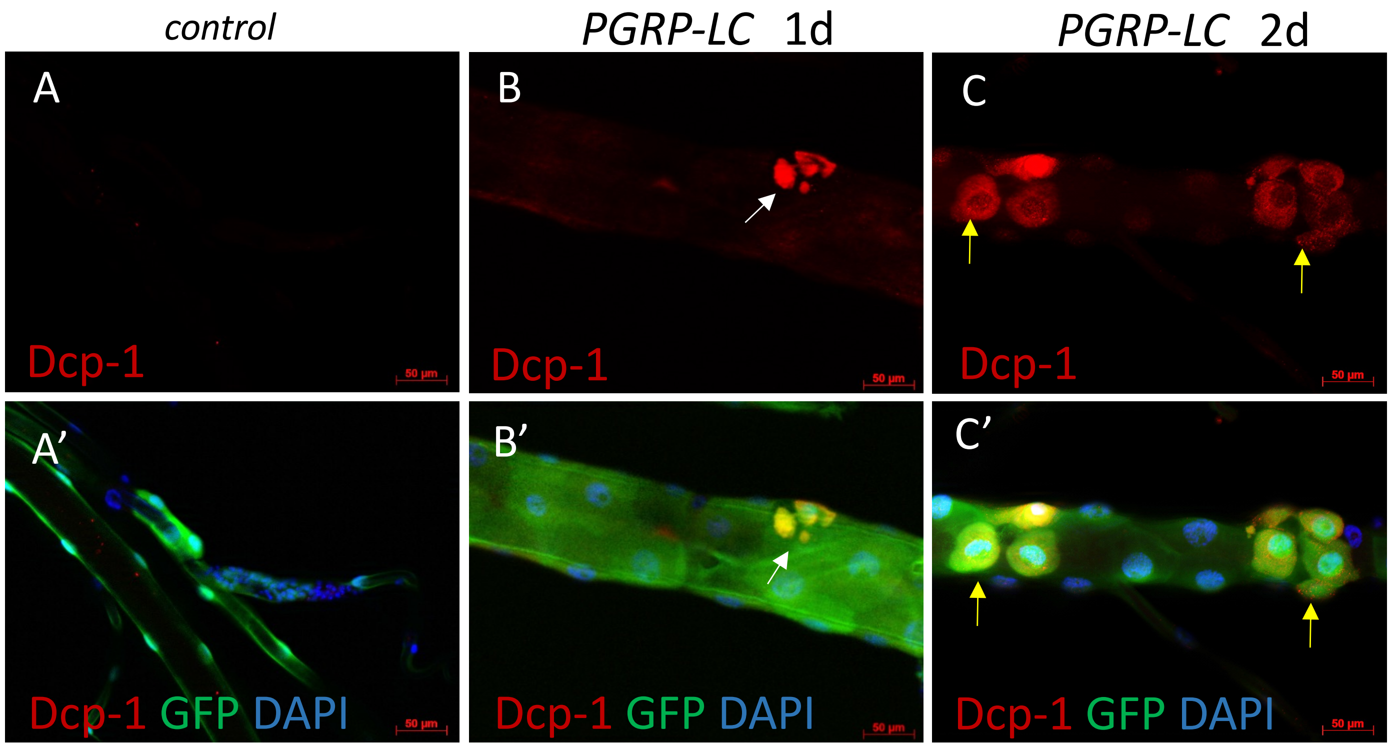


**Supplementary Figure 2. Constitutive activation of IMD signaling in the Drosophila airways leads to apoptosis in the dorsal trunk.** Dissected tracheae from control larvae (A, A'; *btl-Gal4, UAS-GFP; tub-Gal80 [ts]* > $w^{1118}$), PGRP-LC larvae (B, B', C, C'; *btl-Gal4, UAS-GFP; tub-Gal80 [ts]* > *UAS-PGRP-LC(1)*) were stained with Dcp-1 antibody is shown in red. (A'-C') The GFP signal of the trachea is green; the nuclear DAPI signal is in blue. Yellow arrows highlight Dcp-1 positive cells and white arrows show apoptotic vesicles. Animals incubated at 29 °C for 1 d (B, B') or 2 d (C, C') following dissection. Scale bar = 50 μm.

### Supplementary Figure 3

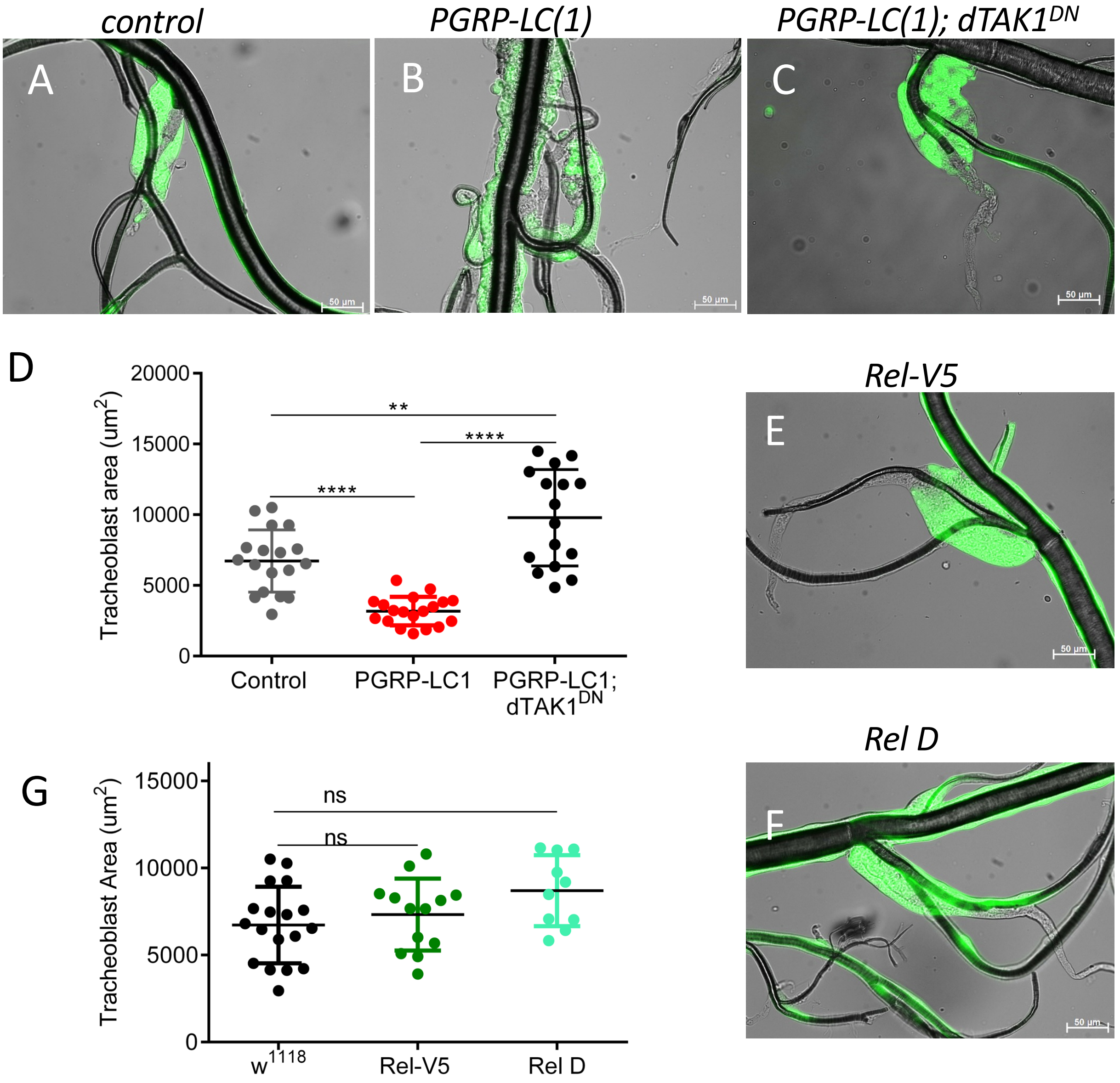


**Supplementary Figure 3. IMD-induced reduction in the tracheoblast area does not depend on the canonical Relish signaling pathway downstream of TAK1.** Dissected tracheae from *btl-Gal4, UAS-GFP; tub-Gal80 [ts] > w$^{1118}$* control larvae (A). Dissected tracheae from *Btl-Gal4, UAS-GFP; tub-Gal80 [ts] > UAS-PGRP-LC(1)* (B). Dissected tracheae from *Btl-Gal4, UAS-GFP; tub-Gal80 [ts] > UAS-PGRP-LC(1); UAS-dTAK1$^{DN}$* (C). (D) Quantitative analysis of the area of T4 tracheoblasts. (E) Dissected tracheae from *btl-Gal4, UAS-GFP; tub-Gal80 [ts] > UAS-Rel-V5*. (F) Dissected tracheae from *btl-Gal4, UAS-GFP; tub-Gal80 [ts] > UAS-Rel-D. (G)* Quantitative analysis of the area of T4 tracheoblasts. Images of control w$^{1118}$ for E and F are the same as A and are not shown again. Scale bar = 50 µm. Statistical significance was evaluated by the Mann–Whitney test, ns, not significant, $**p < 0.01$, $****p < 0.0001$. $n \geq 10$.

#### Supplementary Figure 4

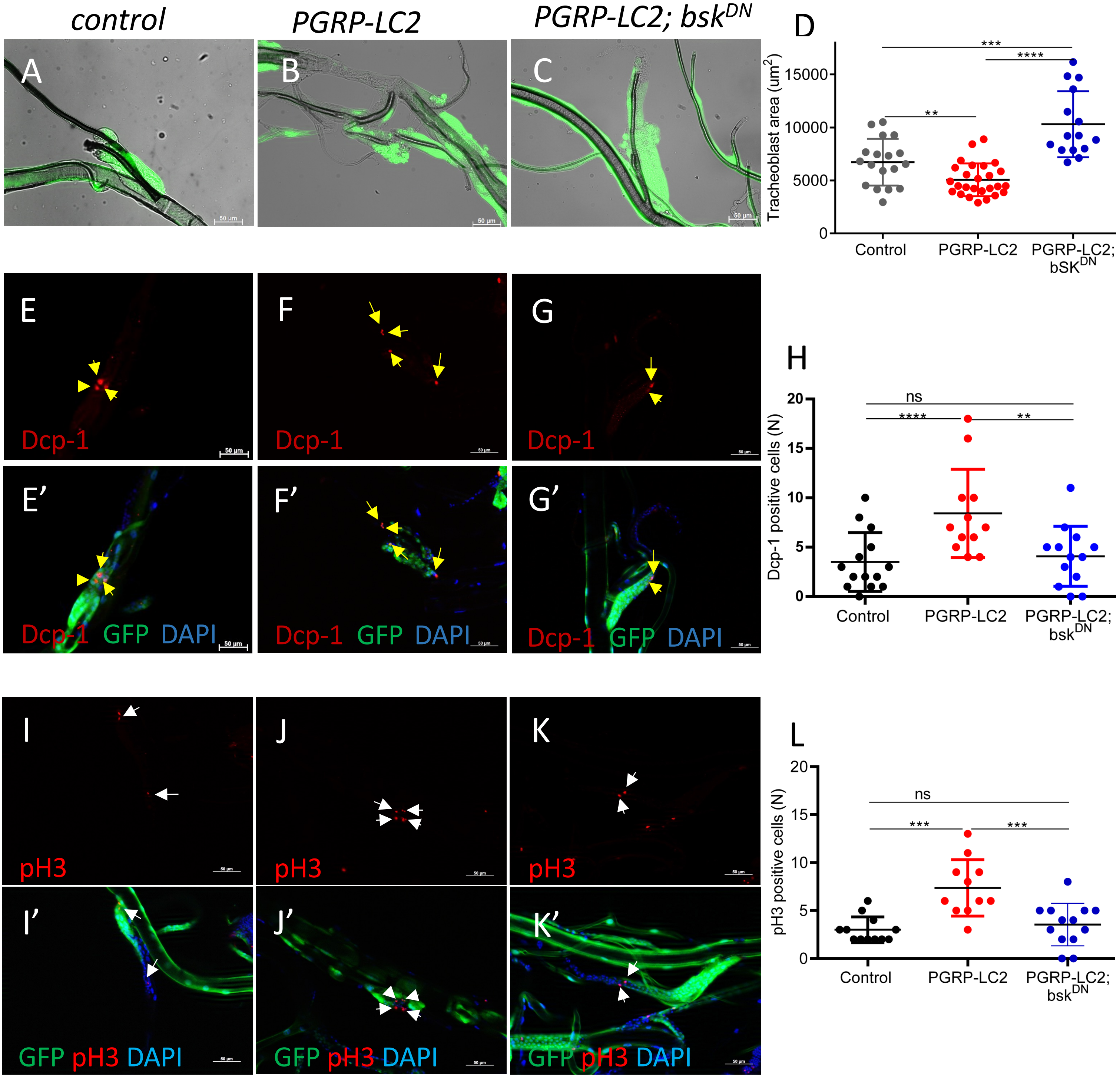


**Supplementary Figure 4. The effects on T4 tracheoblasts caused by activation of IMD signaling are rescued by JNK (bsk) silencing.** (A, E, E' I, I') Dissected tracheae from *btl-Gal4, UAS-GFP; tub-Gal80 [ts] > w$^{1118}$*. (B, F, F', J, J') Tracheae from *Btl-Gal4, UAS-GFP; tub-Gal80 [ts] > UAS-PGRP-LC(2)*. (C, G, G', K, K') Dissected tracheae from *Btl-Gal4, UAS-GFP; tub-Gal80 [ts] > UAS-PGRP-LC(2); UAS-bsk$^{DN}$*. (D) Quantitative analysis of the area of T4 tracheoblasts. (E-H) Effects of PGRP-LC activation and concurrent bsk silencing on Dcp-1-positive cell numbers. (I-L) Effects of PGRP-LC activation and of concurrent bsk silencing on pH3-positive cell numbers. (H) Quantitative analysis of the Dcp-1 positive cells per T4 tracheoblast. (L) Quantitative analysis of the pH3-positive cells per tracheoblast. Arrows mark positive cells. Scale bar = 50 μm. Statistical significance was evaluated by the Mann–Whitney test, ns, not significant, $^{**}p < 0.01$, $^{***}p < 0.001$, $^{****}p < 0.0001$. n≥10.

### Supplementary Figure 5

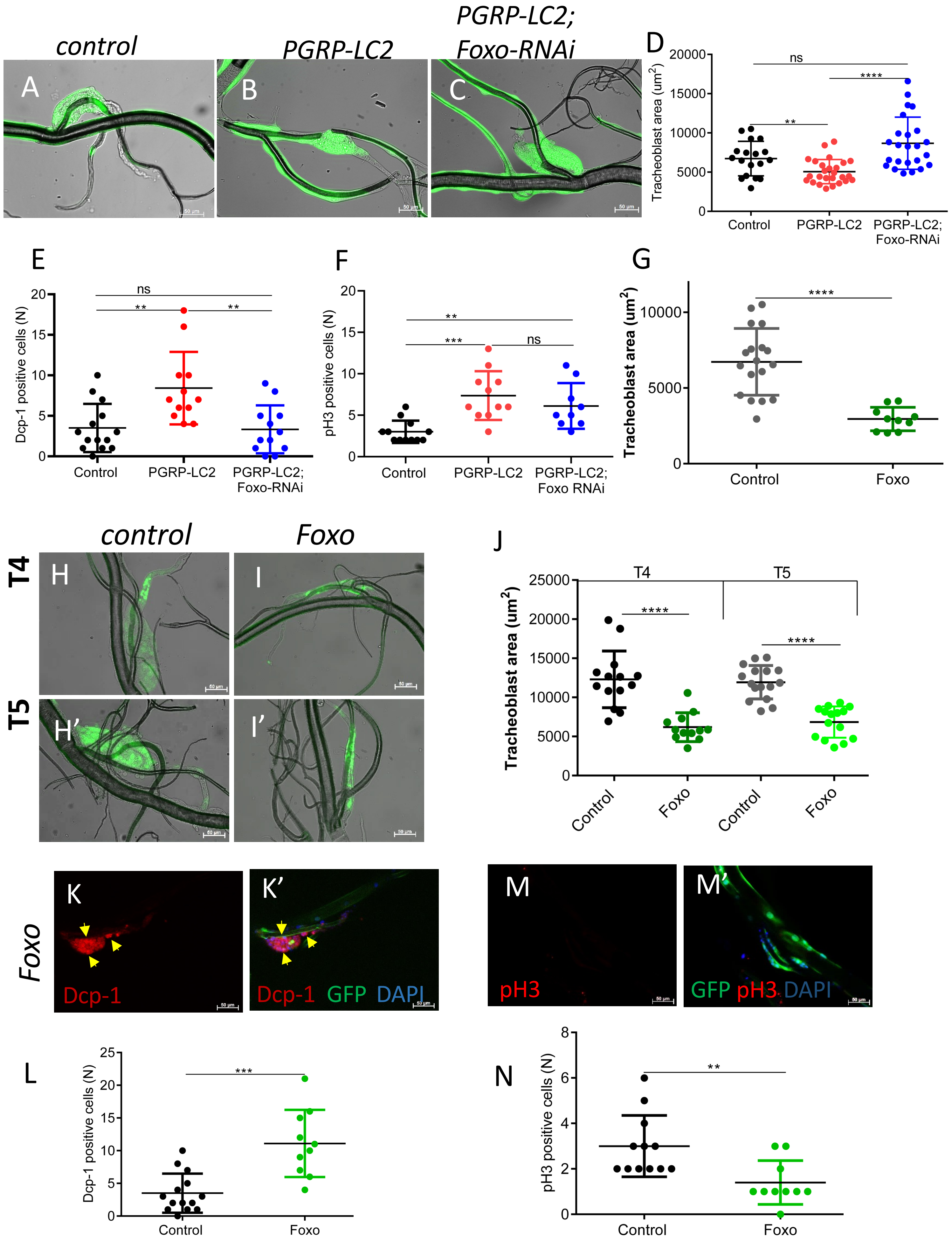

### Supplementary Figure 5

**Supplementary Figure 5. Foxo acts downstream of JNK and is necessary and sufficient for mediating the apoptotic signals in tracheoblasts.** Dissected tracheae from control larvae (A; *btl-Gal4, UAS-GFP; tub-Gal80 [ts] > $w^{1118}$*), of PGRP-LC overexpressing larvae (B; *Btl-Gal4, UAS-GFP; tub-Gal80 [ts] > UAS-PGRP-LC(2)*), those experiencing concurrent expression of *PGRP-LC* and *foxo*-RNAi (C; *Btl-Gal4, UAS-GFP; tub-Gal80 [ts] > UAS-PGRP-LC(2); UAS-foxo-RNAi*), and those overexpression of foxo (K, K', M, M'; *btl-Gal4, UAS-GFP; tub-Gal80 [ts] > UAS-foxo*). Dissected tracheae from control larvae (H, H'; *Ci-gal4, UAS-GFP; gal80 [ts] > $w^{1118}$*), of foxo overexpressing (I, I'; *Ci-gal4, UAS-GFP; gal80 [ts] > UAS-foxo*). (H, I) T4 tracheoblasts. (H', I') T5 tracheoblasts. Anti-dcp1 signals (K, K') and anti-pH3 (M, M') are shown in red, GFP in green, and DAPI in blue. Yellow arrows (pH3 positive cells white arrows) highlight positive cells. Statistical analysis of tracheoblast areas (D, G, J), of dcp1-signals (E, L), and pH3 signals (F, N). Scale bar = 50 μm. Statistical significance was evaluated by the Mann–Whitney test, ns, not significant, ** $p < 0.01$, *** $p < 0.001$, ****$p < 0.0001$, $n \geq 10$.

### Supplementary Figure 6

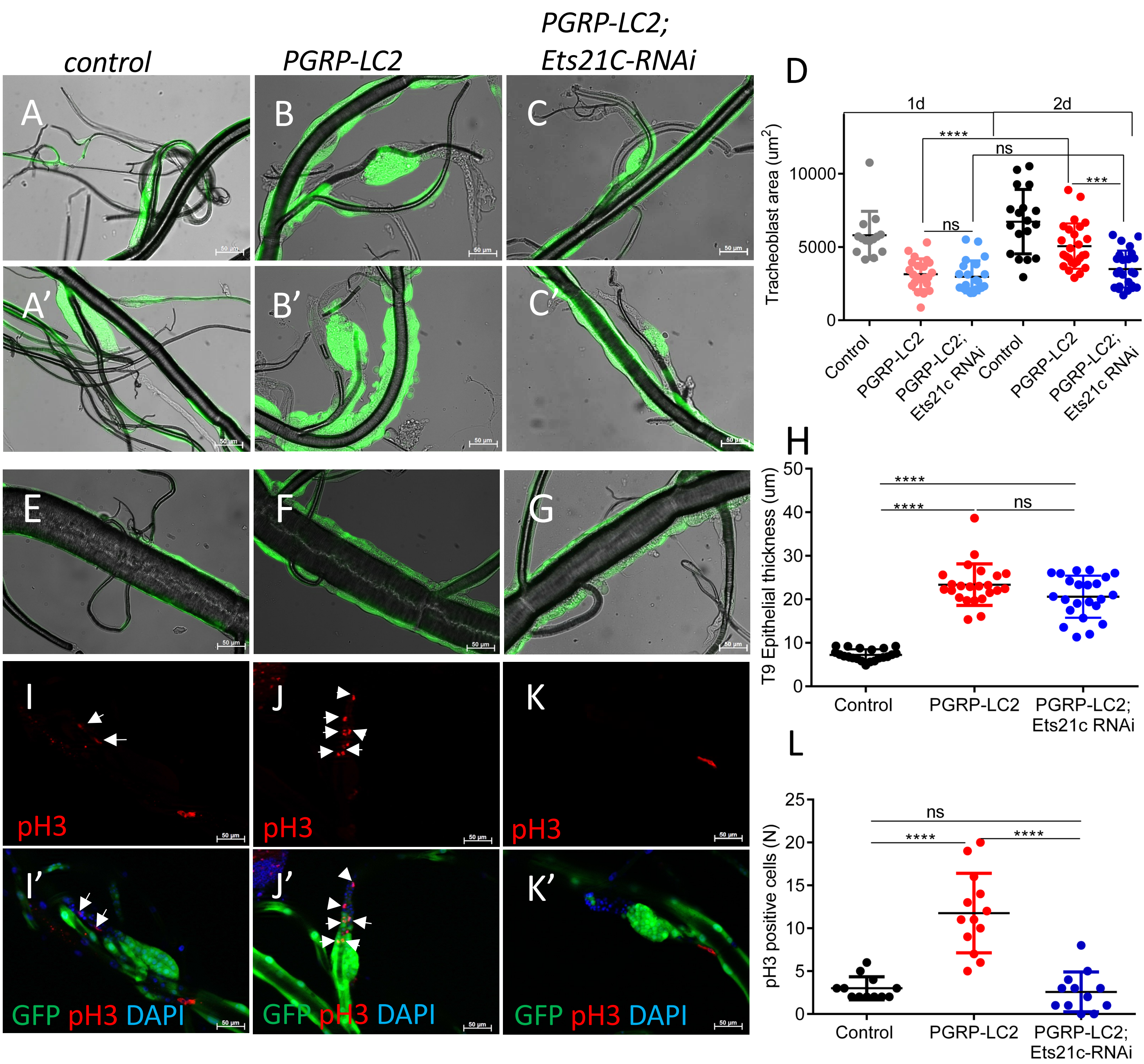


**Supplementary Figure 6. Ets21C enhances the negative effects of PGRP-LC overexpression on tracheoblast size.** Dissected larvae of the control (A. A', E, E', I, I'; *btl-Gal4, UAS-GFP; tub-Gal80[ts]* > *w*$^{1118}$), those experiencing PGRP-LC overexpression (B, B', F, F', J J'; *btl-Gal4, UAS-GFP; tub-Gal80[ts] > UAS-PGRP-LC(2))* and those concurrent expressions of *PGRP-LC* and *Ets21C*-RNAi (C, C', G, G'*; btl-Gal4, UAS-GFP; tub-Gal80[ts] > UAS-PGRP-LC(2), UAS-Ets21C-RNAi).* (A-C) 29°C 1 day. (A'-C', E-G) 29°C 2 day. Analysis of T4 tracheoblast sizes (A-D) with quantitative analyses of tracheoblast sizes (D). Analysis of Tr9 epithelial thickening (E-H) with quantitative analysis of the Tr9 epithelial thickness (H). Anti-pH3 (I-K, I'-M') signals are shown in red, GFP in green and DAPI in blue. White arrows highlight positive cells. Quantitative evaluation of the numbers of pH3 positive cells (L). Scale bar = 50 µm. Statistical significance was evaluated by the Mann–Whitney test, ns, not significant, ***p < 0.001, ****p < 0.0001. n≥10.

### Supplementary Figure 7

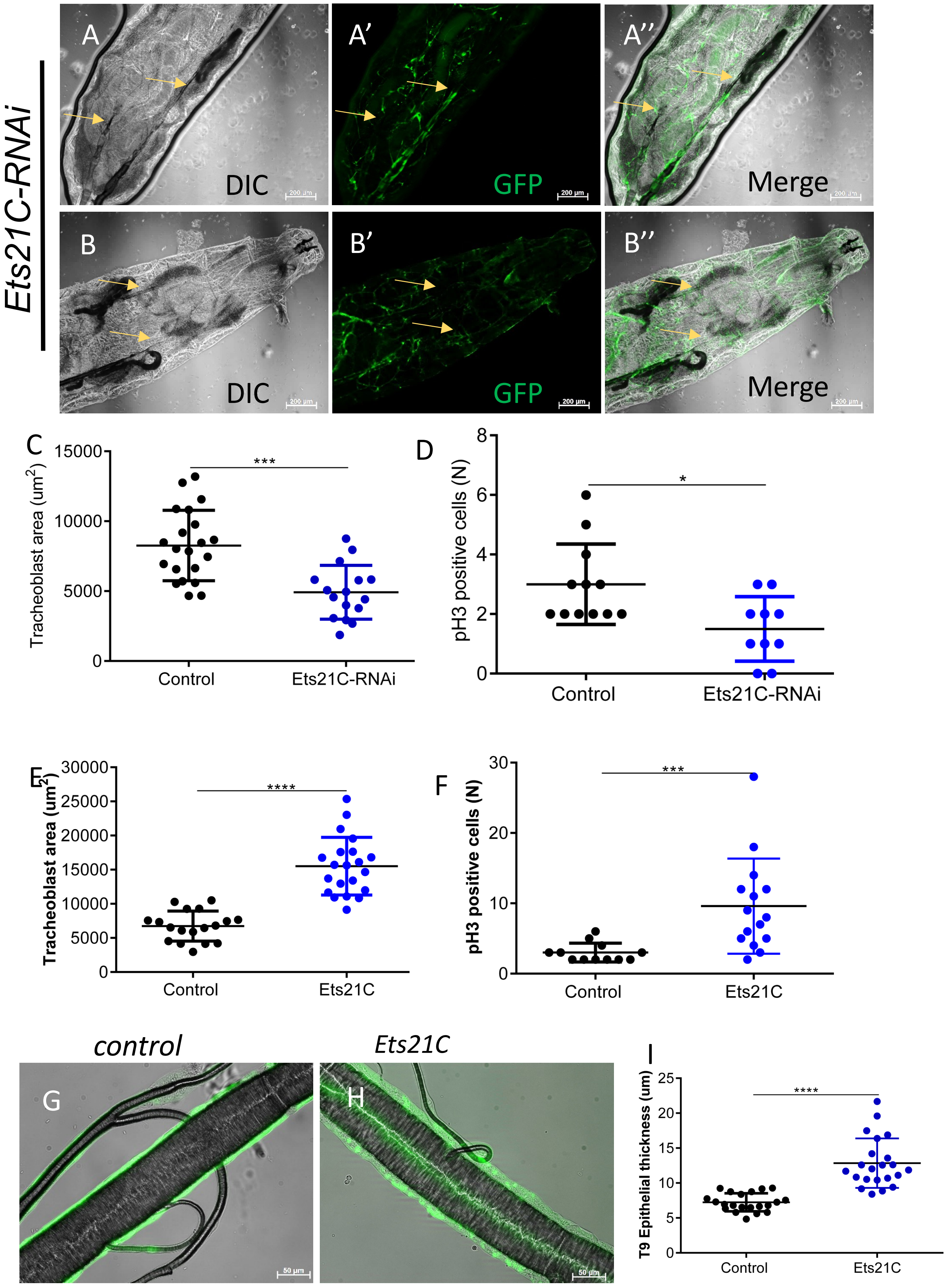

### Supplementary Figure 7

**Supplementary Figure 7. Ets21C is required for larval tracheal development.** (A, A', A'', B, B', B'') Images are from L3 early dying whole larvae that were knocked down Ets21C (*btl-Gal4, UAS-GFP; tub-Gal80 [ts] > UAS-Ets21c RNAi*) in the airway at the embryonic stage. A, A', A'': posterior part of the larva; B, B', B'': anterior part of the larva, the yellow arrows show severe regions of tracheal structural damage. Dissected larval tracheae were isolated from controls (G; *btl-Gal4, UAS-GFP; tub-Gal80 [ts] > $w^{1118}$*) from animals *Ets21C* overexpression (H; *btl-Gal4, UAS-GFP; tub-Gal80 [ts] > UAS-Ets21C*). (I) Quantitative analysis of Tr9 epithelial thickness. Statistical analysis of T4 tracheoblast areas (C, E) and pH3 signals (D, F). (A, A', A'', B, B', B'') scale bar = 200 μm; (G, H,) scale bar = 50 μm. Statistical significance was evaluated by the Mann–Whitney test, *$p < 0.05$, ***$p < 0.001$, ****$p < 0.0001$. $n \geq 10$.

### Supplementary Figure 8

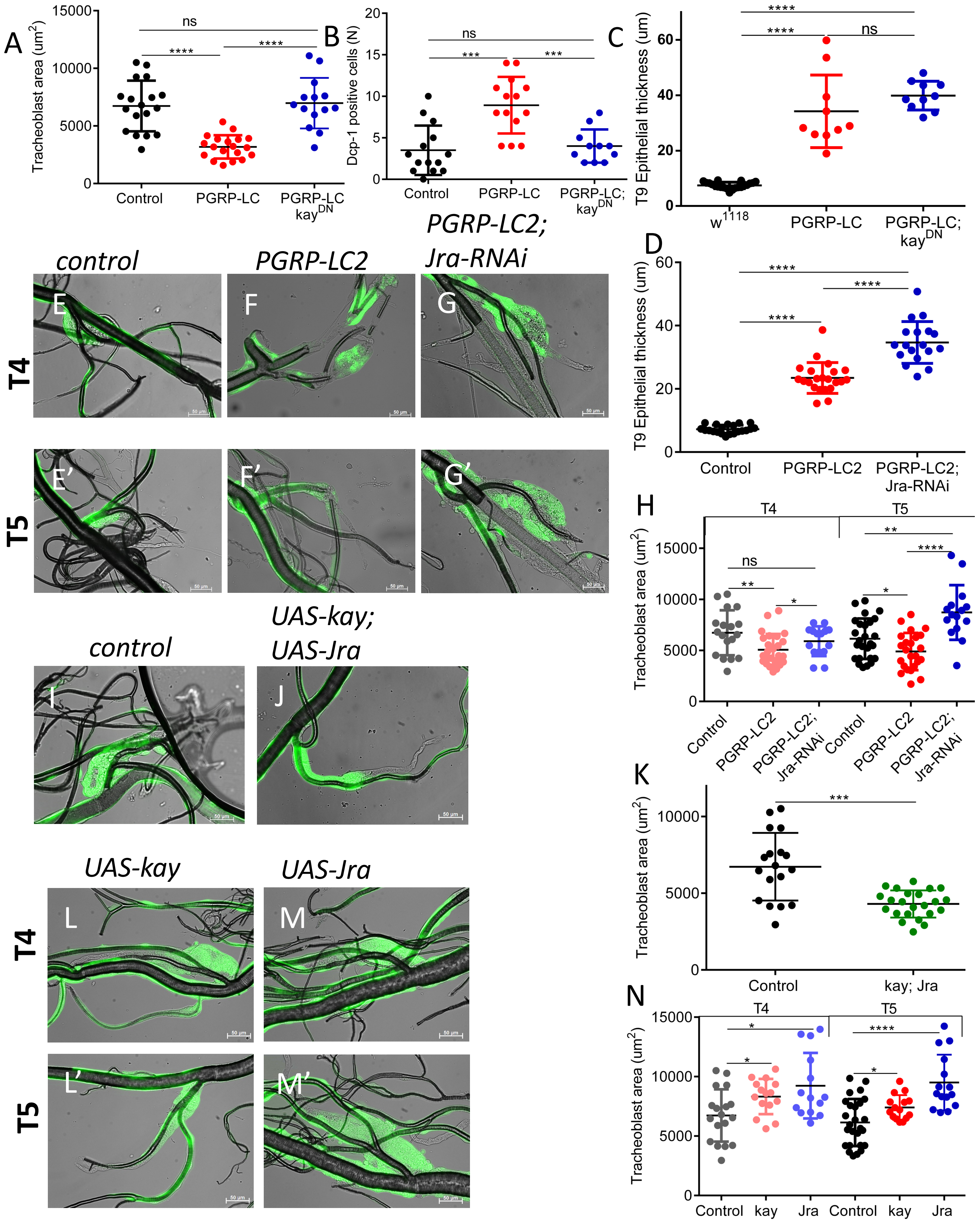

### Supplementary Figure 8

**Supplementary Figure 8. Blocking AP-1-signaling rescues PGRP-LC-induced reduced size of tracheoblasts.**

(A-C) Quantitative analysis of concurrent expression of kay$^{DN}$ and PGRP-LC in the trachea of T4 tracheoblast areas (A), of Dcp1-signals (B), and the Tr9 epithelial thickness (C). Dissected tracheae from *btl-Gal4, UAS-GFP; tub-Gal80 [ts] > $w^{1118}$* control larvae (E, E', I). *Btl-Gal4, UAS-GFP; tub-Gal80 [ts] > UAS-PGRP-LC2* larvae with ectopic expression of PGRP-LC in the trachea (F, F'). *Btl-Gal4, UAS-GFP; tub-Gal80 [ts] > UAS-PGRP-LC2; UAS-Jra-RNAi* larvae with concurrent ectopic expression of PGRP-LC and Jra-RNAi in the trachea (G, G'). *Btl-Gal4, UAS-GFP; tub-Gal80 [ts] > UAS-Kay; UAS-Jra* larvae with concurrent overexpression of kay and Jra in the trachea (I, J). *Btl-Gal4, UAS-GFP; tub-Gal80 [ts] > UAS-Kay* larvae with overexpression of kay in the trachea (L, L') and *btl-Gal4, UAS-GFP; tub-Gal80 [ts] > UAS-Jra* larvae with overexpression of Jra in the trachea (M, M'). (E-G, I, J, L, M) T4 tracheoblasts; (E'-G', L', M') T5 tracheoblasts. Quantitative analysis of concurrent expression of Jra-RNAi and PGRP-LC in the trachea of Tr9 epithelial thickness (D), of T4 and T5 tracheoblast areas (H). (K, N) Statistical analysis of the tracheoblast regions. Scale bar = 50 µm. Statistical significance was evaluated by the Mann–Whitney test, ns, not significant, $*p < 0.05$, $**p < 0.01$, $***p < 0.001$, $****p < 0.0001$. $n \geq 10$.

### Supplementary Figure 9

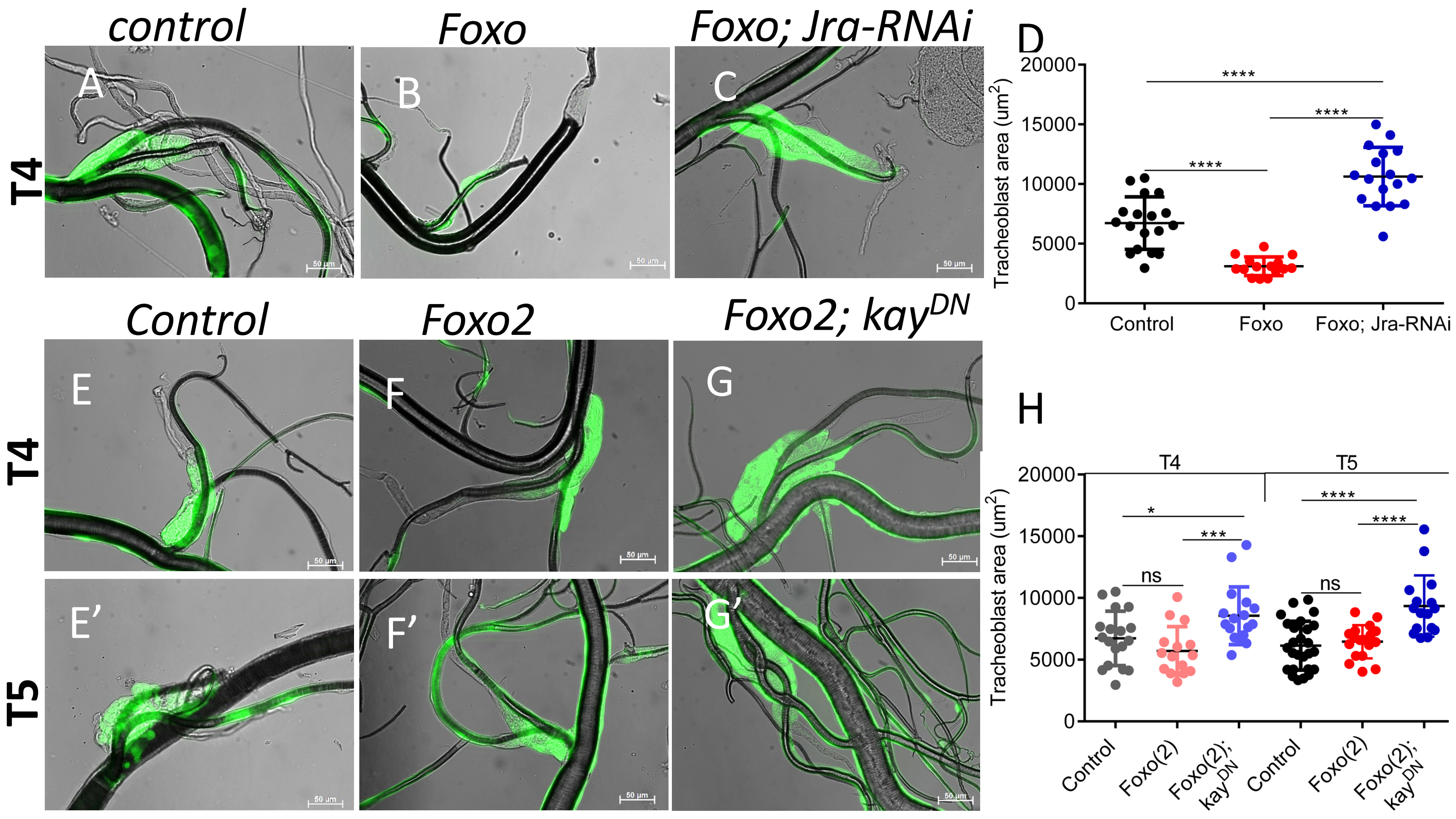


**Supplementary Figure 9. Foxo-induced apoptosis requires the AP1 transcription factor.** Dissected larvae of the control (A, E, E'; *btl-Gal4, UAS-GFP; tub-Gal80[ts] > w[1118]*), those experiencing foxo overexpression (B; *btl-Gal4, UAS-GFP; tub-Gal80[ts] > UAS-foxo* and F, F'; *btl-Gal4, UAS-GFP; tub-Gal80[ts] > UAS-foxo(2)*), those concurrent expressions of *foxo* and *Jra*-RNAi (C*; btl-Gal4, UAS-GFP; tub-Gal80[ts] > UAS-foxo(2), UAS-Jra-RNAi)* and those concurrent expressions of *foxo* and *kay*$^{DN}$ (C, C', G, G'*; btl-Gal4, UAS-GFP; tub-Gal80[ts] > UAS-foxo(2), UAS-kay*$^{DN}$*).* (D) Quantitative analysis of the area of T4 tracheoblasts. Quantitative analysis of concurrent expression of kay$^{DN}$ and foxo in the trachea of T4 and T5 tracheoblast areas (H) Scale bar = 50 µm. Statistical significance was evaluated by the Mann–Whitney test, ns, not significant, *$p < 0.05$, ***$p < 0.001$, ****$p < 0.0001$. n≥10.
